# Adaptive introgression from Pacific herring to Atlantic herring in the brackish Baltic Sea

**DOI:** 10.1101/2025.11.02.685776

**Authors:** Mafalda S. Ferreira, Sabine Felkel, Angela P. Fuentes-Pardo, Jake Goodall, Dandan Wang, Arianna Cocco, Mats Pettersson, Leif Andersson

**Affiliations:** Department of Zoology, Science for Life Laboratory, Stockholm University, Stockholm, Sweden; Department of Medical Biochemistry and Microbiology, Uppsala University, Uppsala, Sweden; Department of Immunology, Genetics and Pathology, Science for Life Laboratory Data Centre, Uppsala University, Uppsala, Sweden; Science for Life Laboratory, Uppsala University, Uppsala, Sweden; Department of Veterinary Integrative Biosciences, Texas A&M University, College Station, USA

## Abstract

The genomic era has revealed that interspecific gene flow can play an important role in adaptive evolution. Here, we report a striking case of adaptive introgression from Pacific into Atlantic herring following the colonization of the brackish Baltic Sea. Gene flow likely occurred in the Arctic, where the distributions of the two species overlap. Although introgression affected only 0.29% of the genome, these regions encompass about 10% of the loci showing strong differentiation between Atlantic and Baltic herring, indicating that natural selection has favored introgressed alleles in functionally important parts of the genome. Adaptive introgression may have been facilitated by shared spawning depths and salinity preferences between Pacific and Baltic herring, in contrast to Atlantic herring, which spawn in deeper, fully marine waters. Introgressed genes such as *THRB* and *SEC16B* may have enhanced visual acuity and lipid metabolism, respectively, promoting adaptation to the brackish Baltic environment.

## Introduction

The ability of species to adapt to new habitats and explore new niches has long fascinated biologists since Darwin. Given the current pace of climate change, understanding the mechanisms of adaptation to new environments has shifted from a matter of curiosity to an imperative for predicting how species may respond to future climates. Recent adaptation to niches opened by deglaciation after the last Glacial Maximum (10,000 years ago) represents a unique opportunity to learn about the evolutionary dynamics and genetic basis of adaptation to new environments.

The slow accumulation of *de novo* mutations makes standing genetic variation a key source for rapid adaptation to new environments ^1^. This is because existing variation is available and has been pre-tested by natural selection. The use of comparative genomics to study adaptation across species has revealed that hybridization between species is an important source of standing genetic variation. Introgressive hybridization has been linked to adaptation in various vertebrate systems, including mammals [e.g. hares ^2,3^], birds [e.g. Darwin’s finches ^4,5^], or fish [e.g. cichlids ^6^]. Importantly, in these examples, introgressive hybridization has allowed adaptation to a novel environment within a matter of decades ^3,5^. Because of the importance of introgressive hybridization for adaptation, understanding how common this is in natural populations is fundamental to determining how species rapidly adapt to new environments.

Atlantic herring (*Clupea harengus*) and Pacific herring (*C. pallasii*) are two sister species that diverged around ∼3 Ma ^7^. The Atlantic and Pacific herring are distributed in the northern parts of the Atlantic and Pacific, respectively (Fig. 1a). During intermediate periods of warming during the last Glaciation ^8,9^, a Pacific herring lineage dispersed through the Arctic and established a population in Europe in the Arctic and Sub-Arctic waters of Russia (the White and Pechora Seas) ^10^. This resulted in a post-glacial contact zone between the two species ^8,11,12^, which has allowed for hybridization to occur between Atlantic and Pacific herring, leading to the origin of a hybrid population in Balsfjord, in Northwestern Norway ^13^. Genomic studies revealed that the Balsfjord population is a Pacific herring population with 25-26% Atlantic herring ancestry and that introgression has likely aided adaptation to the fjord environment. An analysis of the size distribution of introgressed genome fragments in relation to estimated recombination rates suggested that hybridization occurred about 20,000 years ago. The genomic impact of introgression in the Balsfjord population suggests that genetic incompatibilities between Atlantic and Pacific herring might be low, and thus hybridization between the two species could have resulted in introgression in other Atlantic herring populations with an impact on fitness. However, the genomic impact of hybridization across Atlantic herring populations outside Balsfjord has not been investigated.

**Fig. 1.**
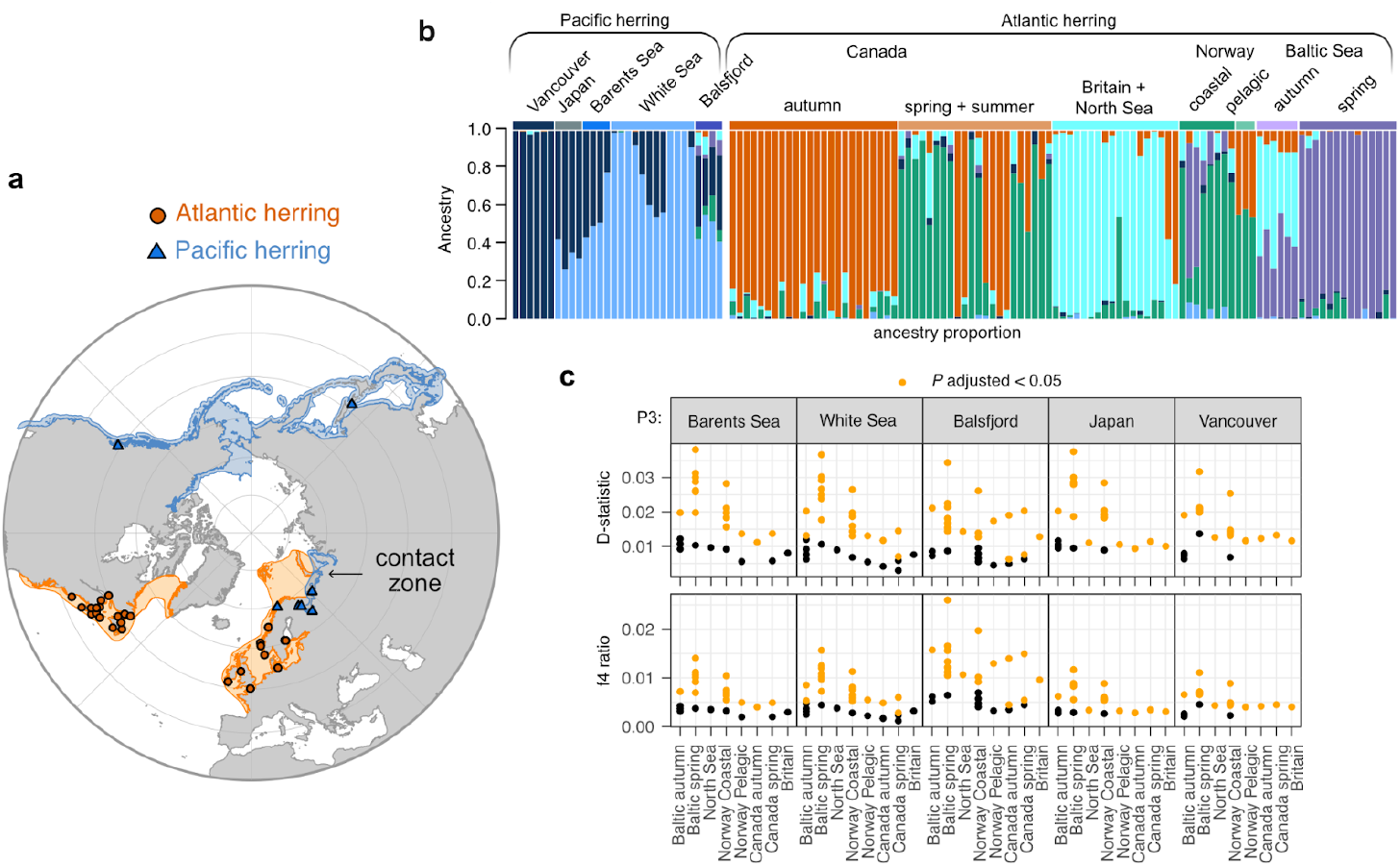
Hybridization history between Atlantic and Arctic Pacific herring. **a** Distribution of Atlantic (orange) and Pacific (blue) herring. Atlantic and Pacific herring individuals included in our dataset are shown as circles and triangles, respectively. **b** Population structure of Atlantic and Pacific herring populations obtained from Admixture (K=6), based on SNPs under selection outside inversion regions. Data based on whole genome sequencing. **c** D-statistic (presence of gene flow) and f4 ratio (admixture proportion) among Atlantic herring populations (P2, or target) and Pacific herring populations (P3, or donor) using European sprat as the outgroup. For each P2, multiple points show results using all other possible Atlantic herring populations as P1. Orange values are significant (P < 0.05).

We combined whole-genome sequencing of Pacific and Atlantic herring populations from across their distributions, encompassing diverse habitats and reproductive strategies, including the Arctic contact zone (Fig. 1a), to investigate the genomic impact of introgression on Atlantic herring. We estimate the proportion of Pacific herring ancestry in Atlantic herring populations. We find that introgression is surprisingly high in spring-spawning Atlantic herring inhabiting the Baltic Sea. We find that introgression of Pacific herring haplotypes occurs in genomic regions overlapping with genes with known functions in visual acuity and body growth, in line with known ecological and phenotypic differences between spawning populations and Baltic and Atlantic herring. Our results provide one of the most striking examples of how inter-species introgression has contributed to genetic adaptation to a novel environment.

## Results

### Genetic structure and admixture of Atlantic and Pacific herring populations

We compiled previously generated whole-genome sequencing data for 95 Atlantic herring individuals representing the distribution of the species in the North Atlantic (Fig. 1a), including the Baltic Sea ^7,13–16^. These individuals represent ecotypes with different spawning seasons (spring, summer, autumn and winter spawning) or ecology (coastal/pelagic, marine/brackish waters). The Pacific herring data (N=30), included samples from a hybrid population in Balsfjord (Norway), the Arctic contact zone with Atlantic herring, Sea of Japan and Vancouver (Fig 1a; Supplementary Table 1). The sequencing depth of coverage of these samples varied between 10X and 48X, with average 27X. All analyses presented are based on this individual whole-genome sequencing data, unless otherwise noted.

We investigated patterns of population genetic structure using Principal Component (PC) analysis (Supplementary Fig. 1) and admixture analysis (Fig. 1b; Supplementary Figs. 2-5). We evaluated results for three SNP datasets, selecting either one SNP every 10 kb along the genome, or using 50 SNPs within regions under selection in herring and excluding or including SNPs overlapping with inversions ^14^. Our results follow previous observations ^13^, with PC1 indicating genetic differentiation between Atlantic and Pacific herring (Supplementary Fig. 1). This separation is also evident in admixture analysis (Fig. 1b; Supplementary Figs 2 to 5). In the PC analysis, while the dataset of windowed SNPs clearly separates the two species, the dataset of SNPs under selection allows finer-scale resolution of population structure within Atlantic herring (Supplementary Figs 1, 3 and 4), as previously shown ^14,15^. As our study focuses on the European contact zone between Pacific and Atlantic herring, we report further results based on the dataset with SNPs under selection outside inversions (Fig. 1b and Supplementary Fig. 5). In the admixture analysis, the best number of ancestral populations (K) was six, according to minimum cross-entropy criterion analysis (Supplementary Fig. 2c). The Balsfjord hybrid population clusters away from other Pacific herring populations, in line with a high proportion of Atlantic herring ancestry (Fig. 1b; Supplementary Figs. 2-5). Within the Atlantic herring, we find individuals grouping by their geographical location and forming clusters corresponding to Canadian, British, Norwegian and Baltic Sea populations, albeit with relatively high levels of admixture between them (Fig. 1b), in line with low levels of genetic differentiation at neutral sites and high migration rates among Atlantic herring populations (F_st_ between 0.01-0.05; N_m_ between 9.9 - 31.7 ^7^). Interestingly, our admixture analysis also suggests Pacific herring ancestry in a few Atlantic herring individuals, particularly the light blue component (see Norway coastal and Baltic sea spring individuals in Fig. 1b) corresponding to Arctic Pacific herring ancestry.

To further explore the impact of gene flow from Pacific to Atlantic herring populations, we calculated genome-wide D-statistics and the proportion of mixing (f4 ratio) between Pacific herring populations (P3) and different Atlantic herring populations (P1 and P2), using the European sprat (*Sprattus sprattus*) as an outgroup (Fig. 1b). We detected significant levels of introgression of Pacific herring alleles into Atlantic herring populations, no matter which Pacific herring population was the putative donor (Fig. 1c). This pattern would be compatible with an ancient admixture event between the two species prior to the divergence of all Atlantic herring populations. The fraction of admixture is clearly the highest in the Baltic Sea spring-spawning herring (Fig. 1c), when considering the White Sea Pacific herring as the donor population. Given the allopatric distribution of these populations, this suggests that Pacific herring alleles occur at higher frequencies in the Baltic Sea spring-spawning herring due to demographic events or selection.

### Distribution and timing of introgression into Baltic spring-spawning herring populations

To locate tracts of Pacific introgression in the Atlantic herring populations, we compared the number of nucleotide differences between a target population and two putative parental populations, in windows across the genome using phased genotype data^13^. This allowed us to classify regions of the genome as high-similarity regions (HSRs) when compared to these parental populations. In our case, HSRs to Pacific herring are likely the result of introgression, as there is no plausible alternative hypothesis. The results indicate low levels of Pacific herring introgression across Atlantic herring populations (Fig. 2a), with values ranging between 0% and 0.29% of the genome. These values are lower than the ones detected with f4-ratios (Fig. 1c) because this analysis requires consistent similarity across the full 20 kb window^13^.

**Fig. 2.**
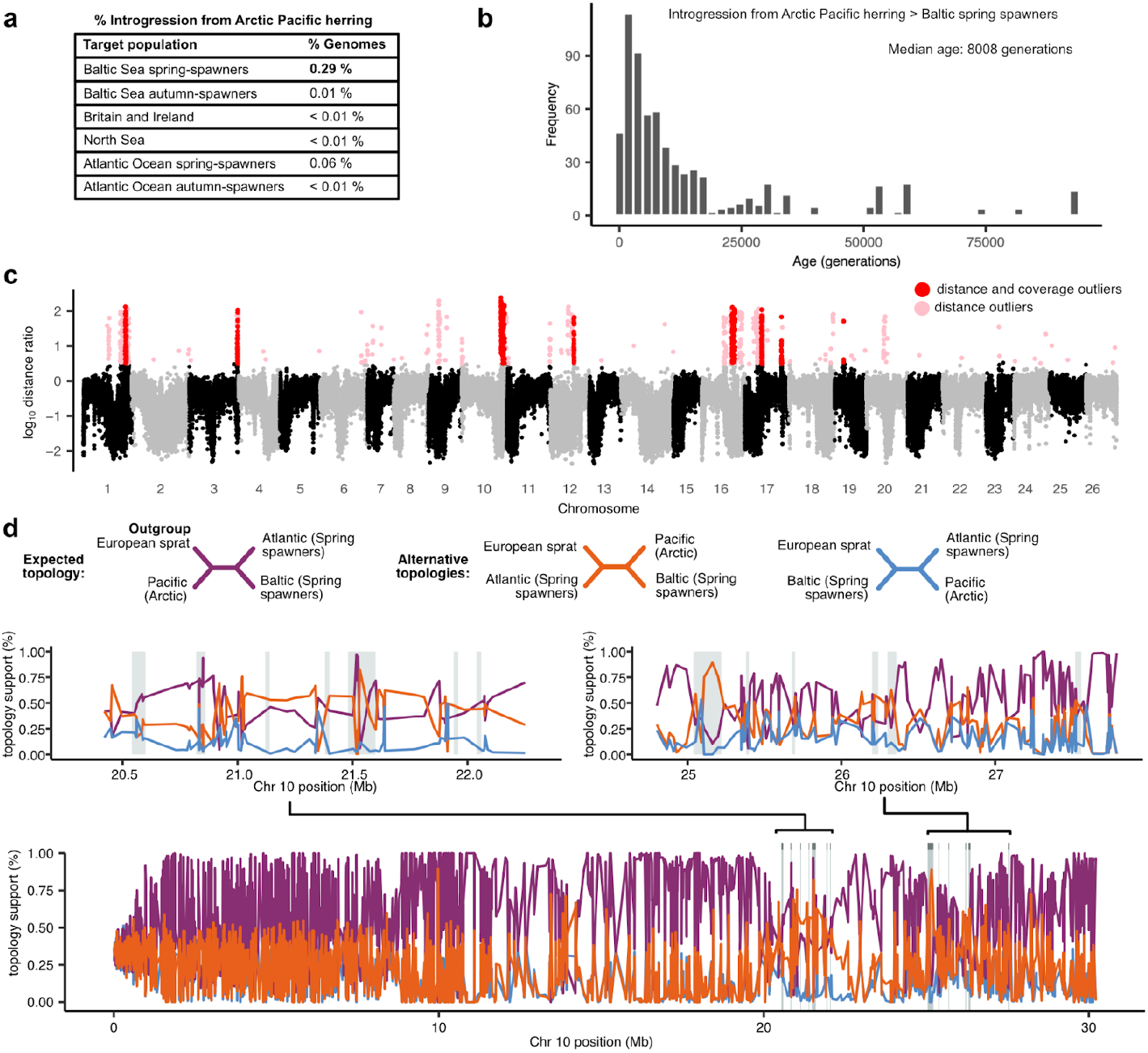
Introgression from Arctic Pacific herring to spring-spawning Baltic herring. **a** Percentage of genomic introgression from Arctic Pacific herring (White Sea + Pechora Sea) to Atlantic herring populations. **b** Distribution of the estimated time since introgression from Arctic Pacific herring to Baltic spring-spawning herring. **c** Genome-wide scan of introgression from Arctic Pacific herring to Baltic spring-spawning Atlantic herring, using a distance-based metric from Pettersson et al. (2023)^13^. Introgression is called (pink or red points) when the genetic distance between individual haplotypes of the target population to the donor population is smaller than the distance to a closely related population. In addition, red points represent windows with support for introgression from more than seven haplotypes. **d** Topology weighting across chromosome 10 (bottom) reveals that introgression regions (gray boxes) show a local increase in a topology supporting introgression between Arctic Pacific herring and spring-spawning Baltic herring (orange alternative topology).

As in the f4-ratio test analysis, the Baltic spring-spawner population stands out again by having higher levels of Arctic Pacific herring introgression (∼0.29% of the genome). Introgression in this population was distributed across 25 major genomic regions and totaled ∼2.09 Mb of the ∼800 Mb herring genome (Fig. 2c; Supplementary Table 3). However, these regions showed strong clustering in the genome, implying that they represent about nine independent regions, some of which have been broken up by recombination since introgression occurred. We used the Atlantic herring recombination map ^17^ and estimated the number of generations of recombination necessary to generate haplotype blocks the same length of the observed introgressed tracks (Fig. 2b). This allowed us to estimate the median age of introgression to be 8,008 generations or 19 KYA [considering a generation time of 2.4 years for Atlantic herring; ^18^], likely in the last Glacial Maximum. Given the possible action of selection by maintaining introgressed blocks in the Baltic spring-spawning population, it is possible that the introgression age is older. In fact, the distribution includes very long blocks of introgression consistent with the action of selection.

We applied independent approaches to confirm HSRs detected in Baltic spring-spawning herring represent introgressed blocks. For instance, the phylogenetic topology in introgressed regions should place Arctic Pacific herring and Baltic spring-spawning herring as sister taxa, contrary to the expected species tree, where Atlantic and Baltic herring populations are more closely related to each other than to Pacific herring populations. Using Twisst, we find that, indeed, the introgressed tree is more represented in HSRs than in the rest of the genome (Fig. 2d and Supplementary Fig. 6). All introgressed regions coincide with regions where we observe a local increase of the introgressed topology, independently of outgroup taxa (either Pacific herring or European sprat (Fig. 2d, Supplementary Figs. 7 and 8). In these regions, we also observe lower genetic differentiation (d_xy;_ Supplementary Fig. 9) between Baltic spring-spawning herring and Arctic Pacific herring than would be expected and significant levels of the fraction of introgression (e.g. f_d_ ^19^ or f_dM_ ^20^), which is expected for introgressed regions (Supplementary Fig. 10).

### Evidence for adaptive introgression

We have previously documented minute genetic differentiation between Atlantic and Baltic herring at selectively neutral loci ^14^. Despite this, F_st_ distribution in this contrast shows a highly significant deviation from the one expected under neutrality with a long tail of SNPs with much higher F_st_ values than expected by mean F_st_ ^16^. Thus, the fact that introgression from Pacific herring was detected primarily in spring-spawning Baltic herring suggests that this represents adaptive introgression. To test this, We used previously published pool-seq data ^13,14,16,21–23^ from 60 populations of Atlantic and Baltic herring to study genetic differentiation in two independent contrasts: (i) Atlantic versus Baltic spring-spawning herring and (ii) Atlantic versus Baltic autumn-spawning herring from Han et al. (2020)^14^. Table 1 lists the most differentiated SNP and their corresponding delta allele frequency (dAF) in each introgressed region for each contrast, together with gene content. The data reveal highly significant genetic differentiation between Atlantic and Baltic herring for all nine major introgressed regions (Table 1). There is also a strong trend that the differentiation at these regions is particularly strong in the contrast for spring-spawners. dAF is higher in Baltic spring-spawners at 18 out of 25 subregions (dAF_Spring_ / dAF_Autumn_ >= 1.2; regions highlighted in blue in Table 1), and dAF is higher in autumn-spawners in one subregion (dAF_Spring_ / dAF_Autumn_ <= 0.8; region Chr12: 16.24-16.34 MB highlighted in orange in Table 1; Supplementary Fig. 11). The results provide strong evidence for adaptive introgression from Pacific to Baltic herring.

**Table 1.**
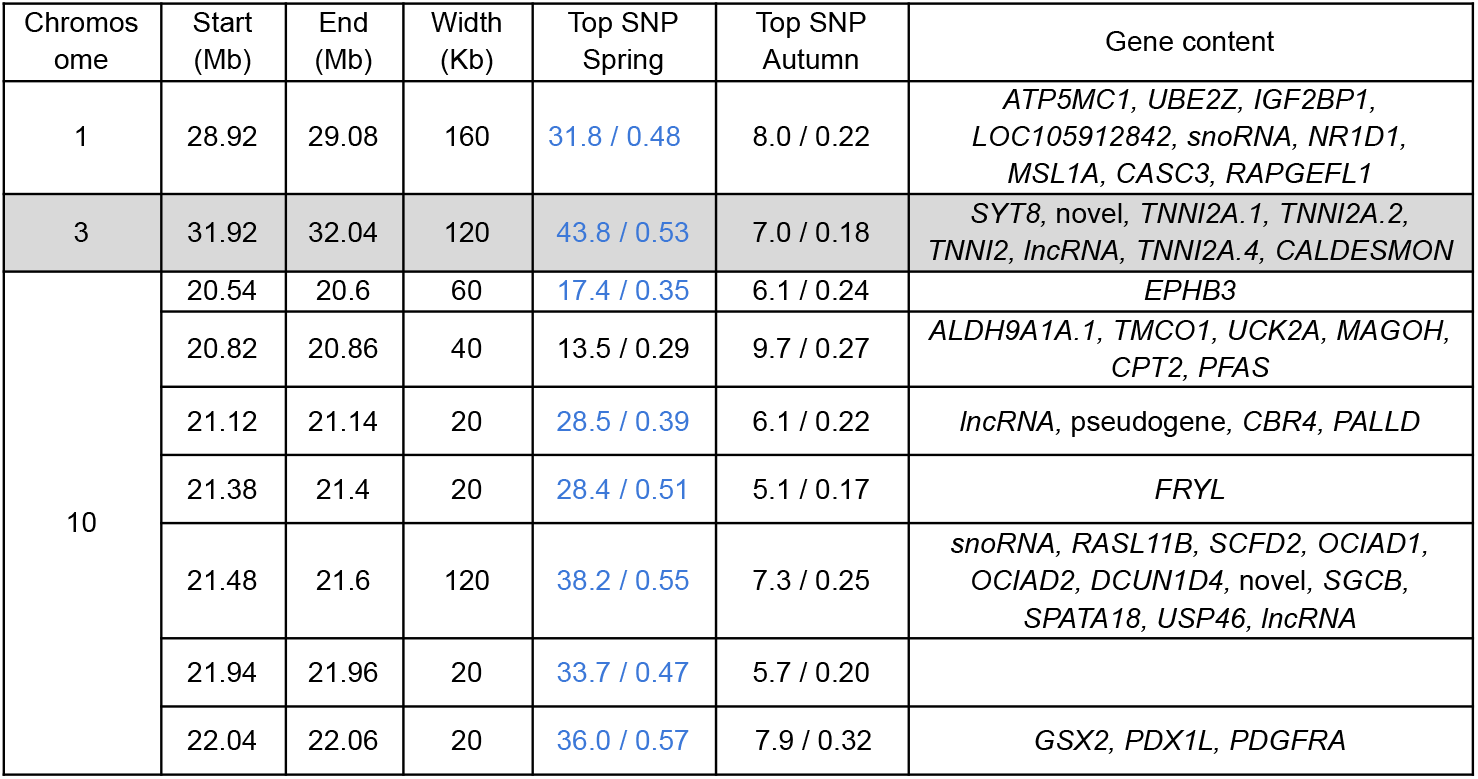
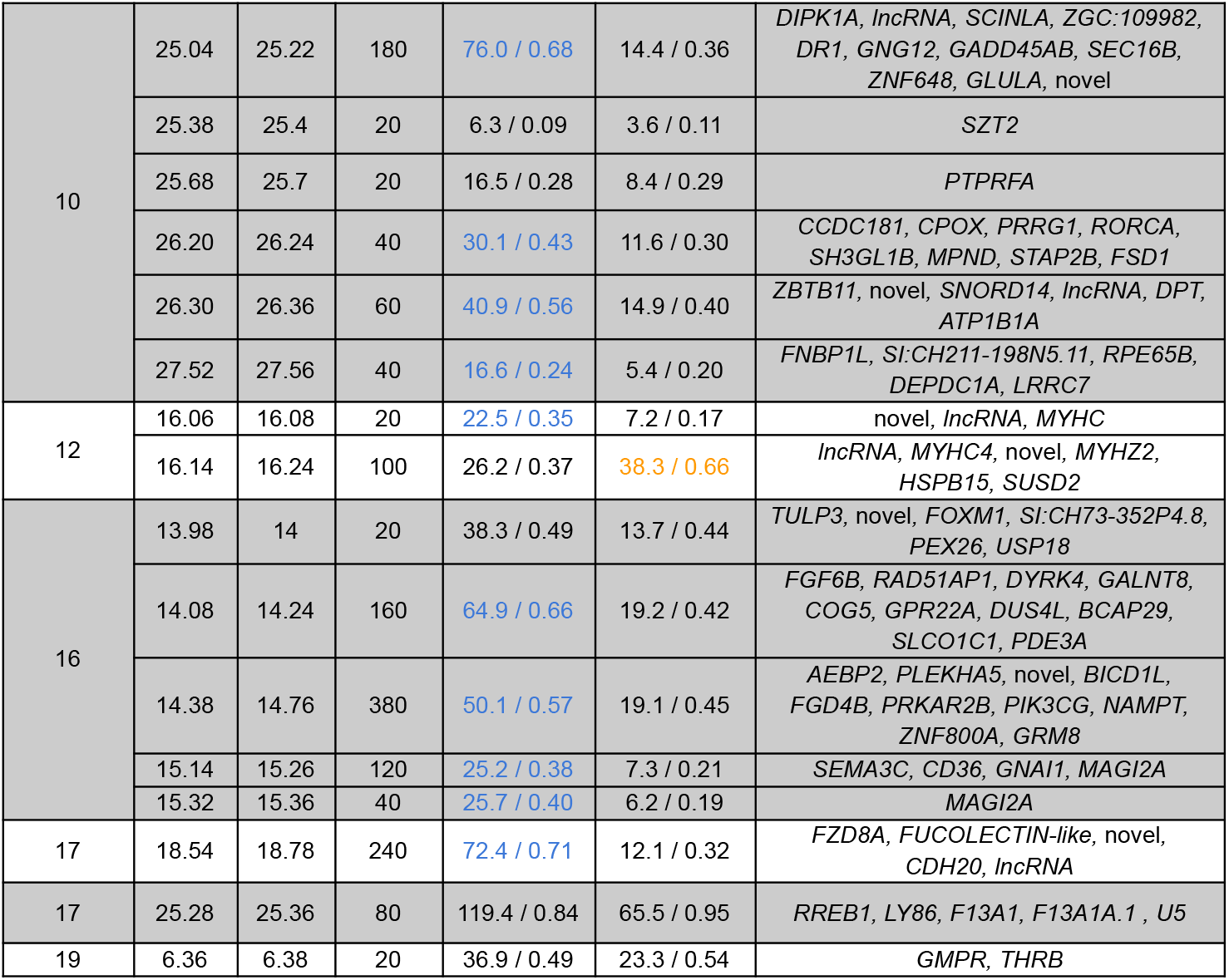
Genomic regions introgressed from Pacific to Baltic herring. For each region we indicate the top differentiated SNP between Atlantic and Baltic within spawning groups (spring or autumn) from Han et al. (2020), (the -log10(*P*) support for genetic differentiation) / (delta allele frequency) and the gene content of the region. The nucleotide positions for top SNPs are in Supplementary Table 3. Regions highlighted in blue and orange are ones where dAF_Spring_ / dAF_Autumn_ >= 1.2 or dAF_Spring_ / dAF_Autumn_ <= 0.8, respectively.

To understand the contribution of introgression for adaptation in Baltic herring, we investigated the gene content of the introgressed regions, which included 166 genes in total on chromosomes 1, 3, 10, 12, 16, 17, and 19 (Table 1; Supplementary Table 4). We performed a Gene Ontology enrichment analysis for these genes, compared to the genomic background (Supplementary Table 5; Supplementary Figure 12). The only two significant terms are the Cellular Component “troponin complex” (FDR adjusted P-value = 3.3 × 10^−18^) and “exon-exon junction complex” (FDR adjusted P-value = 0.004; Supplementary Table 5). The enrichment of the “troponin complex” should be explained by a cluster of troponin I2 genes present in the introgressed region in chromosome 3 (Supplementary Table 4).

We investigated the potential functional significance of SNPs within introgression regions by selecting SNPs that are simultaneously differentiated between Atlantic and Baltic spring-spawners but not differentiated between Baltic spring-spawners and Arctic Pacific herring (significance established as -log_10_ < 6.4 after Bonferroni multiple test correction assuming 137,098 SNPs within introgression regions; Supplementary Table 6). We found a total of 3,535 SNPs following this pattern. Most of them (3,352 SNPs) were annotated as having a “modifier” function and overlap with intergenic, intronic, or putative regulatory regions. Then, 96 mutations are classified as likely having low impact, as they are synonymous mutations, 84 moderate impact missense mutations, and three high impact missense mutations (Supplementary Table 6). These three high-impact mutations are located in the genes *IGF2BP1* (splice donor variant, chromosome 1), *PDGFRA* (splice acceptor variant, chromosome 10) and *SLCO1C1* (splice donor variant, chromosome 16).

The 25 subregions of introgression (Table 1) involved in the adaptation of spring-spawning herring to the Baltic Sea are expected to be differentially introgressed and under positive selection in spring-spawning herring when compared to autumn-spawners. We thus inspected signatures of divergence within regions of introgression comparing Baltic spring- and autumn-spawners and Arctic Pacific herring (Supplementary Fig. 13). We see that divergence between Baltic spring-spawners and Arctic Pacific herring is below background in introgressed regions, while divergence between autumn-spawners and White Sea herring in the same regions is within the expected range (Supplementary Fig. 13a). Importantly, divergence between autumn- and spring-spawners from the Baltic Sea, as expected, is above genomic background in introgressed regions (Supplementary Fig. 13a). At the same time, most introgressed regions show significant signatures of recent positive selection in Baltic spring-spawning herring when compared with autumn-spawners (standard xpEHH < -2: Supplementary Fig. 13b). Nevertheless, this trend is not consistent for all introgressed regions. Of the 2.09 Mb of introgressed regions identified (Fig. 2c), about 37% (780 kb) show simultaneously low d_xy_ between spring-spawners and Arctic Pacific herring and high d_xy_ between autumn-spawners and White Sea (Supplementary Fig. 13a, Supplementary Table 7). Of these, 340 kb (16% of total introgressed regions) show also a signature of recent positive selection in Baltic spring-spawners when compared to Baltic autumn-spawners (Supplementary Fig. 13b, Supplementary Table 7). We consider the evidence for adaptive introgression for these regions totalling 340 kb of the genome particularly strong (Supplementary Table 7) and we further investigate the contribution of these regions to adaptation in the sections below.

### Adaptive introgression overlaps with non-synonymous substitutions in SEC16B

One of the most striking subregions of introgression is on chromosome 10, a 180 Kb region between positions 25.04 Mb and 25.22 Mb (Fig. 3; Supplementary Table 7). This region shows a clear sign of discordance with the expected topology (Fig. 2d), where Baltic spring-spawners are placed as sister taxa to Arctic Pacific herring (Fig. 3d), and is under positive selection in Baltic spring-spawners when compared to Baltic autumn-spawners (Fig. 3c). We again used pool-seq data to study the allele frequency differences across this region (Fig. 3a,b). In this region, we find strong genetic differentiation between Baltic spring- and autumn-spawners, with the highest values of differentiation clustering around the *SEC16B* and *ZNF648* genes. Interestingly, the most common allele in Baltic spring-spawners and Arctic Pacific herring is rare in all Atlantic herring populations except in some Norwegian fjord populations, where it reaches intermediate frequencies (Fig. 3b). This suggests that the introgressed Pacific herring alleles segregate in some populations of Atlantic herring outside the Baltic Sea. It should be noted that salinity is often lower in Norwegian fjords compared with the open ocean, particularly in spring due to the influx of meltwater.

**Fig. 3.**
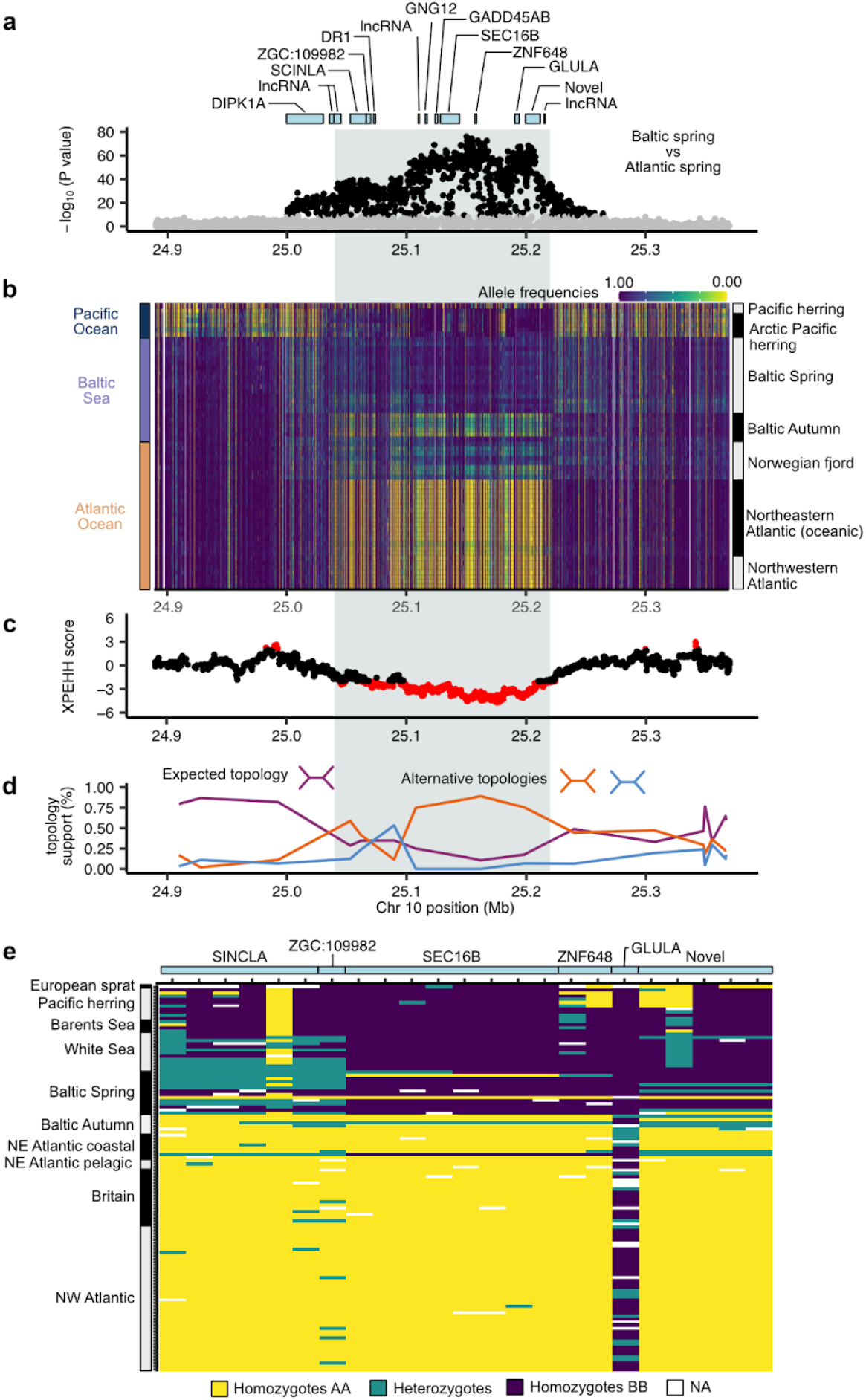
Adaptive introgression at the *SEC16B* locus in spring-spawning Baltic herring. **a** Allele frequency differentiation between spring-spawning Baltic herring and spring-spawning Atlantic herring in the Chr10:25.04-25.22 Mb region. *P*-values for each SNP were calculated with a chi-square test from allele count data from pool-seq data ^14,15^. **b** Allele frequency heat map for Atlantic and Pacific herring populations within the introgressed region. **c** Cross population extended haplotype homozygosity comparing spring- and autumn-spawning Baltic herring. Negative values suggest positive selection in spring-spawners. Red values are 2 standard deviations from the mean. **d** Topology weighting calculated with Twisst for the three expected topologies relating European sprat (outgroup), White Sea Pacific herring, Baltic spring-spawning herring and Atlantic spring-spawning herring (see Fig. 2 for trees). **e** Genotype heatmap for non-synonymous substitutions for genes within the introgression region. In all plots, the introgression region is symbolized as a grey box.

We investigated the potential functional significance of highly differentiated SNPs within these genes. Remarkably, we find as many as eight non-synonymous substitutions in *SEC16B* and two in *ZNF648* (Fig. 3e) with strong allele frequency differences between Baltic spring-spawning herring and other Atlantic herring populations. Genotype data derived from individual whole genome sequencing show that most Baltic spring-spawners are homozygous for the Pacific allele. This pattern of segregation is partly repeated in other non-synonymous mutations, including those in *ZNF648*.

We used the sequence from the closely related European sprat, which separated about twelve million years ago from the common ancestor of Atlantic and Pacific herring ^7^, as a reference to deduce if the Atlantic or Pacific variant represented the ancestral or derived state. Surprisingly, the Atlantic variant is the derived state at all eight non-synonymous substitutions in *SEC16B* (Fig. 3e). The large number of non-synonymous mutations in *SEC16B* in the Atlantic herring populations (except Baltic spring-spawners) relative to Pacific herring and European sprat suggests accelerated protein evolution. We used codeml (PAML) to test for different scenarios of branch selection along a tree, including *SEC16B* sequences from Atlantic and Pacific herring and three outgroups (European sprat, rainbow trout, and zebrafish), and found that all models allowing for accelerated evolution in Atlantic spring-spawning herring are significant (Supplementary Fig. 14 and 15). In a scenario where both Baltic and Atlantic spring-spawning herring branches evolve under selection, the results suggest that the Pacific herring allele has evolved under purifying selection (dN/dS ≈ 0), whereas the Atlantic spring-spawning allele shows a dN/dS ≈ 1.7, indicating positive selection. (Supplementary Fig. 14).

### Adaptive introgression in THRB important for visual acuity in the Baltic Sea

Another striking signal of interspecies adaptive introgression occurs in a narrow region on chromosome 19: 6.36 to 6.38 Mb (Fig. 4; Supplementary Table 7). Analysis of pool-seq data shows highly significant genetic differentiation between spring-spawning Atlantic and Baltic herring in this region (Fig. 4a). Introgressed alleles from Arctic Pacific herring occur at a particularly high frequency in spring-spawning Baltic herring, but occurs at a lower frequency for instance in autumn-spawning Baltic herring and Atlantic herring fjord populations (Fig. 4b).

**Fig. 4.**
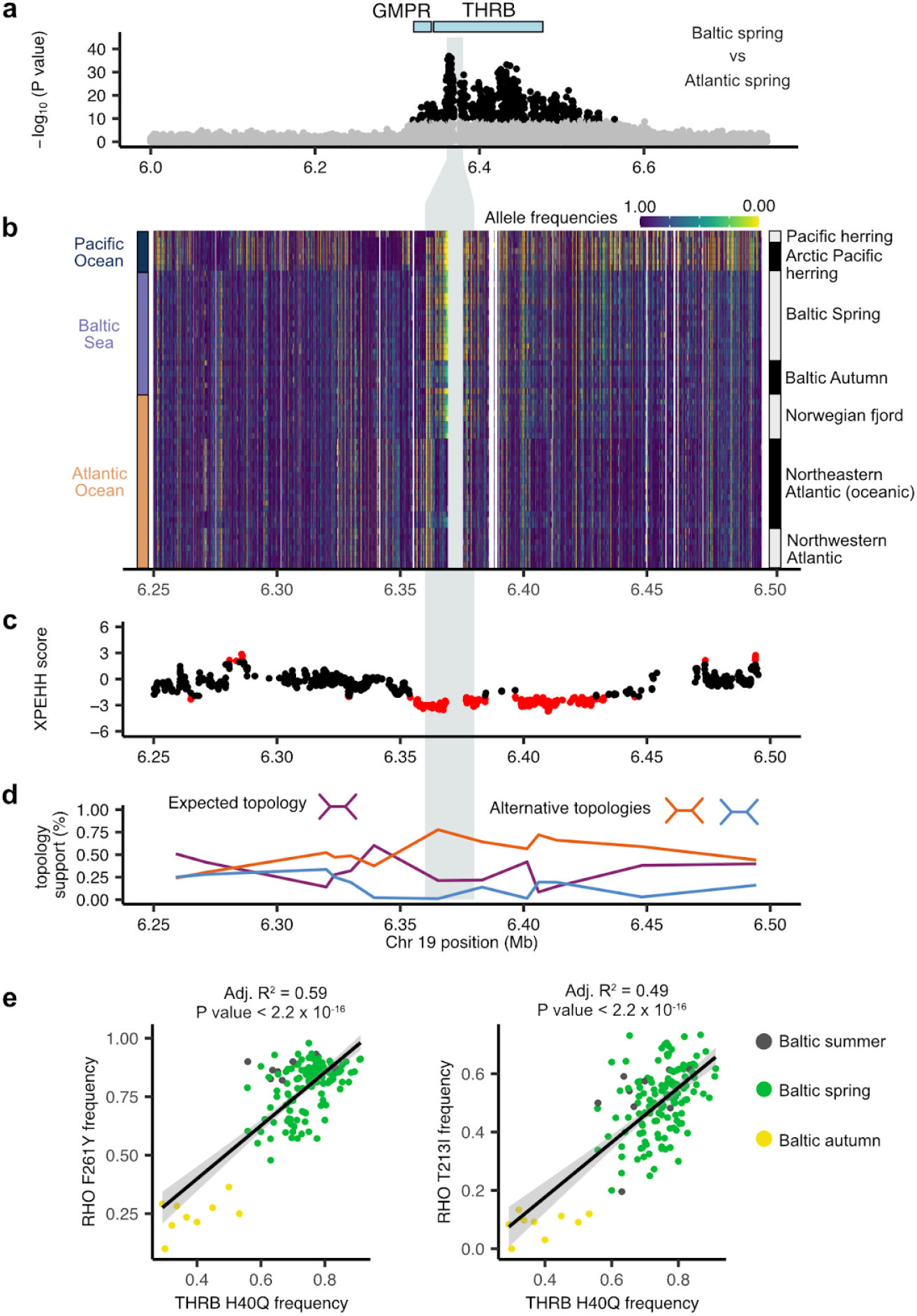
Adaptive introgression in *THRB*. **a** Allele frequency differentiation between spring-spawning Baltic herring and spring-spawning Atlantic herring in the Chr19:6.36-6.38 Mb region. *P*-values for each SNP were calculated with a chi-square test from allele count data from pool-seq data ^14^. **b** Allele frequency heat map for Atlantic and Pacific herring populations within the introgressed region and flanking areas. **c** Cross population extended haplotype homozygosity comparing spring- and autumn-spawning Baltic herring. Negative values suggest positive selection in spring-spawners. Red values are 2 standard deviations from the mean. **d** Topology weighting calculated with Twisst for the three expected topologies relating European sprat (outgroup), White Sea Pacific herring, Baltic spring-spawning herring, and Atlantic spring-spawning herring (see Fig. 2 for trees). **e** Allele frequency correlation between H40Q *THRB* mutation and two Rhodopsin mutations in Baltic herring. Allele frequency data is derived from SNP-chip data from ^28^ for 4,678 spawning fish.

There is also evidence for selection acting on that region in spring-spawning Baltic herring (Fig. 4c) and Twisst analysis support the alternative topology for this region with spring-spawning Baltic herring being sister to Pacific herring from the Arctic Sea (Fig. 4d). The introgression region overlaps a single gene, *THRB* (thyroid hormone receptor beta), and is close to *GMPR* (guanosine monophosphate reductase), located just outside the peak. An examination of the coding sequence of these genes identified a missense mutation H40Q located in the first exon of one *THRB* isoform (THRB-203 based on ENSEMBL 109 Atlantic herring annotation; see Materials and Methods; Supplementary Table 8, Supplementary Fig. 16 and 17a). The frequency of the variant allele (40Q) shows a very striking pattern (Supplementary Fig. 17b). All Atlantic and Pacific populations spawning in marine environments (35-36 PSU) are fixed for the Atlantic allele (40H) whereas the Arctic Pacific allele (40Q) occurs at a very high frequency in spring-spawning Baltic herring. Interestingly, populations spawning at intermediate salinities (fjord populations, Skagerrak, Kattegat and the transition zone between Atlantic Ocean and the Baltic Sea) as well as autumn-spawning Baltic herring show low to intermediate frequencies of the Pacific allele (Supplementary Fig. 17). Furthermore, the H40 allele, associated with the marine environment, is also the ancestral allele shared with European sprat (Supplementary Fig. 17c). The results for Atlantic herring spawning in Ringköbing Fjord and Landvik are particularly informative because these populations spawn in brackish water, similar to Baltic populations, but feed as adults in fully marine conditions. This suggests that selection at this locus is driven by environmental conditions during spawning and early development rather than by conditions experienced during adulthood. The strong differentiation between spring- and autumn-spawning Baltic herring further indicates that the selective factor is not salinity itself, since both groups experience the same salinity during spawning, but rather an environmental factor correlated with salinity.

*THRB* is a highly pleiotropic gene with important functions in many cell types and contributes to metabolic regulation ^24^. Interestingly, the missense mutation occurs in the first exon of a transcript isoform denoted *THRB2*, primarily expressed in the retina ^24^. Furthermore, work in zebrafish has demonstrated that *THRB* plays a critical role in the development of red cone photoreceptors and expression of long-wave opsins (LWS1 and LWS2) needed for the detection of long-wave visual light ^25–27^. We have previously identified two missense mutations (F261Y and I213T) in herring *rhodopsin* (*RHO*) that have contributed to genetic adaptation to the red-shifted light conditions in the brackish Baltic Sea ^23^. We took advantage of data from a recent comprehensive genetic screen of Baltic herring populations ^28^ to explore genetic correlation of allele frequencies between the *THRB* missense mutation from chromosome 19 and the two *RHO* missense mutations from chromosome 4 (Fig. 4e). There is a very strong and highly significant correlation in allele frequencies between the two loci and the difference between spring- and autumn-spawning herring is consistently replicated. The result suggests that the *THRB* polymorphism may reflect natural selection on visual acuity in the Baltic Sea.

Previous work on zebrafish ^25–27^ shows that *THRB* regulates the expression of long-wave opsins (LWS1 and LWS2). The fact that the Arctic Pacific allele (40Q) is not completely fixed in spring-spawning Baltic herring provided an opportunity to test the hypothesis that *THRB* is affecting vision by regulating the relative proportion of red cone photoreceptors and/or the expression of LWS1 and/or LWS2. Quantitative RT-PCR analysis revealed low expression of *LWS2* in both genotypes (Supplementary Table 9 and 10) but a significant upregulation of *LWS1* in *Q/Q* homozygotes compared with *Q/H* heterozygotes (Supplementary Fig. 18). This result strongly supports our interpretation that the introgression of the Arctic Pacific *THRB* allele has contributed to adaptation to the red-shifted light conditions in the brackish Baltic Sea, and we thus consider the *THRB* missense mutation as a candidate causal mutation.

## Discussion

We provide clear evidence of interspecies introgression from Arctic Pacific into Baltic herring, and show that this introgression has contributed to adaptation to the brackish Baltic Sea. This is reflected in the strong genetic differentiation between Atlantic and Baltic herring at introgressed regions (Table 1). Our results thus suggest that roughly 10% of all loci showing high differentiation between Atlantic and Baltic herring populations (Han et al. 2020), despite minimal divergence at neutral sites, can be attributed to adaptive introgression from Pacific herring. The sister species Atlantic and Pacific herring diverged from a common ancestor about three million years ago ^7^, and were likely completely isolated during glacial periods. However, gene flow would have been possible during interglacial periods, such as the present day, when the Arctic Sea is ice-free during summer and supports plankton production.

The two sister species occupy similar ecological roles as key links between plankton production and higher trophic levels, yet they differ in spawning behaviour. Pacific herring spawn in shallow water, such as estuaries with lower salinity, and deposit their eggs on seaweed, whereas Atlantic herring spawn in deeper water and lay eggs on bedrocks ^29^. Interestingly, the spring-spawning Baltic herring is more akin to that of the Pacific herring, with spawning taking place in shallow water, where eggs are often deposited on seaweed ^30^. Thus, it is possible that introgression of gene variants from Pacific to Baltic herring can be adaptive.

Introgression from Atlantic herring to Pacific herring, resulting in a hybrid population in a Norwegian subarctic fjord, has been well documented ^11–13^. Here, we provide evidence of introgression in the opposite direction. Our results indicate that Pacific herring from the Arctic Sea acted as the donor population and that gene flow likely occurred in the contact zone between Atlantic and Pacific herring (Fig. 1a). Because both species are broadcast spawners that reproduce in large aggregations where eggs and sperm are released without mate choice, strong prezygotic reproductive isolation is unlikely to have evolved. This lack of isolation may reflect their enormous population sizes, which reduce the likelihood that genetic incompatibilities arise through drift. Moreover, selection to avoid hybridization is expected to be weak in broadcast spawners, since the individual cost of occasional hybrid fertilizations is low when each female releases tens of thousands of eggs. Under such conditions, postzygotic selection likely plays a key role in both purging deleterious introgressed alleles and in retaining adaptive ones. This interpretation is supported by our results: the few introgressed regions (0.29% of the genome) are concentrated in genomic regions under selection among locally adapted Atlantic herring populations (Table 1).

We identified nine major regions of adaptive introgression from Pacific to Baltic herring (Table 1). Some of these have been broken up by recombination, resulting in 25 subregions with clear evidence for introgression (Table 1). The majority of subregions show genetic differentiation predominantly or exclusively in spring-spawning populations. One of these loci is located on chromosome 19 and includes a single gene, *THRB* (Fig. 4). Interestingly, strong genetic differentiation at this locus between Atlantic and Baltic populations is also observed in the closely related European sprat ^31^. Our data suggests that this is related to vision and photoreceptor expression. We identified a missense mutation H40Q in the THRB2 transcript expressed in the retina, and observed significant differences in long-wave opsin (*LWS1*) expression between *THRB* genotypes in spring-spawning Baltic herring (Supplementary Fig. 17). *LWS1* is highly expressed in red cone photoreceptors, and zebrafish knockouts show that *THRB* regulates *LWS1* expression ^27^. Light conditions in brackish and freshwater environments are red-shifted compared to marine environments due to absorption of blue wavelengths by dissolved organic matter ^32^. Thus, the upregulation of LWS1 expression and possibly increased red cone photoreceptor abundance in herring carrying the introgressed *THRB* allele would increase visual acuity in the brackish Baltic environment. This is consistent with the correlation of the introgressed THRB allele with the missense mutation Phe261Tyr in *rhodopsin* (*RHO*) causing a red-shifted light absorbance of about 10 nm ^23^. This *RHO* mutation is present in one third of freshwater and brackish water fish species while rare in marine fish ^23^. Our results combined with evidence from European sprat suggest that *THRB* may play a similar adaptive role to varying light conditions. The introgressed *THRB* His40Gln missense mutation occurs at ∼ 0% in marine Atlantic herring, ∼ 50% in autumn-spawning Baltic herring, and ∼ 85% in spring-spawning Baltic herring (Supplementary Fig. 16), a pattern also observed for the RHO Phe261Tyr missense mutation (Fig. 4e). A plausible explanation for this allele frequency difference between autumn- and spring-spawners is that selection is strongest during the juvenile stage, when the larvae need good vision to find food and avoid predation. Seasonal differences in light conditions are due to river runoffs of meltwater in the spring ^23^. This interpretation is consistent with the difference in spawning behaviour between Atlantic and Pacific herring, as Pacific herring often spawns in estuaries, where larvae experience turbid, red-shifted light, whereas Atlantic herring spawn in deeper, clearer waters.

The most prominent signal of adaptive introgression was detected on chromosome 10, spanning a 2.5 Mb region (25.04 and 27.56 Mb; Table 1). This large block likely contains multiple genes with functional significance. The strongest peak of differentiation between Atlantic and Baltic herring occurs within a subregion (Chr10:25.04-25.22 Mb) overlapping *SEC16B*, where Pacific and Atlantic alleles differ by eight missense mutations (Fig. 3). All missense mutations are derived in the Atlantic allele and are under accelerated protein evolution, indicating that the ancestral *SEC16B* protein sequence was reintroduced into Baltic herring through introgression from the Pacific lineage.

*SEC16B* encodes a protein involved in intracellular lipid trafficking ^33^, and in mammals it has been associated with variation in lipid absorption ^34^, body composition and metabolic traits such as obesity ^35,36^ and type II diabetes ^37^. These findings suggest that the introgressed *SEC16B* allele may contribute to lipid metabolism in Baltic herring, a critical trait in the energy-limited brackish environment of the Baltic Sea. Baltic herring, especially spring-spawners inhabiting northern parts of the Baltic Sea, are smaller and leaner than Atlantic herring, likely reflecting lower plankton productivity in the Baltic Sea ^38,39^. Thus, Pacific-derived alleles at *SEC16B* may enhance energy storage and reproductive success under these conditions.

Notably, the Pacific *SEC16B* allele is more frequent in spring-spawning than in autumn-spawning Baltic herring (Table 1), consistent with differences in energy demands between these reproductive strategies ^40,41^. Autumn-spawning larvae grow very slowly during winter but are ready to take advantage of plankton production in the spring. In contrast, spring-spawning herring have to starve throughout the winter and be ready to spawn in the spring when plankton production starts, which the larvae need for feeding. Summer-spawning Pacific herring from the Arctic Sea experience a similar energetic constraint as spring-spawning Baltic herring, which may explain why the Pacific *SEC16B* allele is particularly advantageous in this ecotype. Our interpretation that adaptive introgression of *SEC16B* relates to metabolic regulation is consistent with the marked increase in spring-spawning Baltic herring during the 20th century, likely driven by eutrophication and enhanced plankton production that favored this reproductive strategy ^42^. Further experimental studies are required to fully understand the functional significance of the introgression of the Pacific *SEC16B* allele to Baltic herring.

The Atlantic herring is now added to a rather long list of examples of introgression between taxa, including the ones between archaic and modern humans ^43^, Darwin’s finches ^44^, cichlids ^6^, and different species of hares ^45^. The herring stands out by the relatively deep split between the species involved [about three million years ^7^] and by the large number of loci with conclusive evidence for adaptive introgression. The herring, therefore, provides an outstanding example of how low frequency, introgressed variants present in an ancestral population become part of the standing genetic variation that contributes to rapid genetic adaptation after colonization of a new environment. The fact that the Atlantic herring has, within the last 8,000 years, colonized the Baltic Sea with its striking differences in both abiotic conditions, such as salinity, and biotic conditions (such as plankton production and paucity of large carnivores), makes it an excellent model for genetic studies of ecological adaptation.

## Materials & Methods

### Processing of individual whole-genome re-sequencing data

We combined whole genome sequencing data from 65 Atlantic herring and 30 Pacific herring individuals ^7,13–16^ mapped to the Atlantic herring reference genome ^17^. We performed genotype calling for each sample using Haplotyper and joint calling with GenotyperGVCFs within the Sentieon wrapper (release 202112.07; ^46^), which implements GATK4 HaplotypeCaller ^47^. We used GATK 3.4.0 and bcftools 1.17 ^48^ to filter genotypes. We filtered indels and genotypes with RMSMappingQuality lower than 40.0, MQRankSum lower than −12.5, ReadPosRankSum lower than −8.0, QualByDepth lower than 2.0, FisherStrand higher than 60.0 and StrandOddsRatio lower than 3.0. We further filtered variants with genotype quality below 20, read depth below 2 or higher than 3 times the average depth of the sample. We removed positions where genotypes were missing for more than 20% of the individuals, and filtered variant positions where minor allele frequency was below 5%. Our final dataset comprised 23,223,645 positions, of which 1,352,818 were SNPs. We annotated variant positions using snpEff (v 5.1d) with OracleJDK 11.0.9 ^49^. We also performed statistical phasing of the variants using Beagle (v5.4)^50^.

### Processing of European sprat long-read data

We used available long-read data from the outgroup species ^31^, the European sprat, to map and call genotypes to use in our analysis. We mapped long-read data for one European sprat individual using minimap2 (v2.26) using assembly-to-assembly alignment preset option “asm20” ^51^, and applied the same genotype calling approach combining Sentieon and GATK described above to call and filter variant and invariant sites.

### Processing of Atlantic and Pacific herring pool-sequencing data

We also include pool-seq data from 60 populations to infer allele frequencies ^13,14,16,21–23^. We obtained allele counts for each pool of individuals already corrected for raw read depth [N_eff_ correction; ^15,52^]. For chi-square calculations between Atlantic Ocean spring-spawners, Baltic Sea spring- and autumn-spawners, we used population contrasts defined by Han et al. (2020)^14^. We kept biallelic positions with minor allele frequency above 0.01, following Han et al. (2020).

### Population structure analysis

To conduct population structure analysis with high-coverage individual whole-genome sequencing data, we construct three different SNP datasets. Based on best practices for population structure analysis, the first dataset consisted of unlinked variant sites that were distanced at least 10 kb from each other (total of 67,566 SNPs). Based on previous results suggesting that Atlantic herring populations are also well distinguished by strongly differentiated SNPs ^14^, we selected the 50 most differentiated SNPs within genomic regions putatively under selection from Han et al. (2020)^14^ resulting in 15,231 SNPs. From these we also generated a third dataset where we excluded SNPs within previously identified inversion regions ^7,14^, resulting in 11,492 SNPs. For each analysis described below, we repeated the analysis with each SNP dataset and including or not Pacific herring individuals, which allowed us to detect different aspects of population structure in Atlantic herring (see Results section). We conducted a Principal Component Analysis using PLINK (v1.90b4.9 ^53^) using default settings. We also conducted an Ancestry Admixture analysis using sNMF (v1.2) ^54^. We repeated the sNMF analysis for values of K from 2 to 10. For each K, the analysis was repeated across five random initializations (different seeds: 10, 20, 30, 40, 50), to check if results were qualitatively consistent. As results were similar among runs, we analyzed the results of seed 10. We used cross-entropy analysis to estimate the best value of K as the one where the error is minimized (in this case, K=6).

### Estimating admixture among populations and species

We used Dsuite (v0.4 r43) ^55^ to estimate levels of ancestral hybridization and admixture among populations. We used Dtrios to calculate genome-wide estimates of D (ABBA-BABA) and f4-ratio statistics for all possible trios of populations and species, using the European sprat as the outgroup and a jackknife block size of 1000 SNPs. As we wanted to test for admixture between Pacific herring populations and the relationship between the sister species is known ^7^, we analyzed results where the P3 population were Pacific herring populations (i.e, Vancouver, Japanese, White Sea, Barents Sea or Balsfjord), and P2 and P1 were all possible pairs of Atlantic herring populations.

We used Dinvestigate to calculate the proportion of introgression (f_dM_ ^20^) in windows of 50 SNPs with a step of 25 SNPs along the genome. Here, we considered P3 as White Sea or Barent Sea Pacific herring, and P2 as Baltic spring-spawning or Baltic autumn-spawning herring. We used Canada spring-spawners as P1.

### Detecting putative regions of adaptive introgression

To detect introgression regions, we followed the approach described and developed by Pettersson et al (2023) ^13^ to detect introgression in Atlantic herring. Briefly, we identified highly supported regions (HSRs) of introgression using a metric based on the relative Hamming distances to the nearest Atlantic and Pacific reference haplotypes. The reference populations for each species varied depending on the target population (see Supplementary Table 2). The genome was divided into 20 kb windows, and within each window, reference haplotypes were extracted from predefined comparison groups, according to Supplementary Table 2. For each haplotype, we calculated the minimum Hamming distance to the reference haplotypes in both groups. Windows with very low SNP diversity (maximum distance <20) were excluded to avoid spurious signals. Then, to generate an introgression score, we calculated the ratio of these distances per window. This ratio normalizes for variation in SNP density and is symmetric in log space, facilitating detection of introgressed regions in either direction. We considered introgressed regions where at least 7 haplotypes passed the threshold. Finally, introgressed regions located within 50 kb of each other were merged into contiguous blocks, resulting in 25 subregions of introgression which we summarize in 9 regions (Table 1). All computations were performed using custom R scripts adapted from the *HaploDistScan* R package (https://github.com/HaploDistScan/HaploDistScan). We annotated introgression regions using the function *makeTxDbFromBiomart()* from the GenomicFeatures R package (v 1.58; ^56^) to retrieve Atlantic herring annotations from ENSEMBL (109). Then we determined the gene content of introgressed regions and flanking 20 kb regions using *subsetByOverlaps()* from the package GenomicRanges [v. 1.58; ^56^)].

To support the results of the introgression scan, we further calculated summary statistics conventionally used to detect and/or support introgression in the genome. We calculated two measures of topology discordance (topology weighting and f_dM_) and genetic divergence (d_xy_). We calculated f_dM_ using Dsuite as described above (see ***Estimating admixture among populations and species***). To estimate topology weighting we used Twisst ^57^ (accessed from https://github.com/simonhmartin/twisst on February 2025) using Atlantic herring phased data (see ***Processing of individual whole-genome re-sequencing data***), and European sprat genotypes as outgroup. We used *phyml_sliding_windows*.*py* to generate neighbor joining trees (--optimise n) in windows of 100 genotype sites (--windType sites, -w 100) with a minimum of 10 non-missing genotypes per individual (-Mi 10). We then used *twisst*.*py* to perform topology weighting for a group of four taxa: Canada spring-spawners, Baltic Sea spring-spawners, Baltic Sea autumn-spawners, Arctic Pacific herring and the outgroup European sprat (see Supplementary Table 1 for the groupings) using the default “complete” method.

We calculated d_xy_ between Baltic Sea spring-spawners, Baltic Sea autumn-spawners, and Arctic Pacific herring populations (as defined in Supplementary Table 1) in 20-kb windows across the genome using *pixy* (v.1.2.5.beta1; ^58^). For these initial calculations, all Baltic spring-spawning individuals were included, regardless of their introgression genotype. The resulting values (Supplementary Fig. 9) provided a first assessment of how well the introgression scan detects introgression, by comparing d_xy_ in introgressed regions against the genomic background. However, Arctic Pacific herring alleles are not fully fixed in Baltic spring-spawning populations (see Figs. 3b and 4b). As a result, calculating d_xy_ between all Baltic spring-spawners and Arctic Pacific herring does not necessarily yield the expected signal of reduced divergence in introgressed regions ^59^. To better evaluate the potential adaptive role of introgression, we therefore focused on regions showing strong evidence of introgression, where divergence between Arctic Pacific herring and Baltic spring-spawners is particularly low. For this refined analysis, we re-calculated d_xy_ using only Baltic spring-spawners that were homozygous for Arctic Pacific herring alleles. To identify these individuals, we used the Hamming distances from the introgression scan: individuals with both haplotypes unusually close to Arctic Pacific herring were considered likely homozygotes (custom script: *find_homozygote_indv_dxy*.*R* available in https://github.com/MafaldaSFerreira/Baltic_herr_introgression). For each introgressed region, we then calculated d_xy_ between these homozygotes and Arctic Pacific herring using *pixy*. These results are shown in Supplementary Fig. 13.

We further consider putative regions of adaptive introgression in Baltic spring-spawning herring ones where we see a strong signature of positive selection in Baltic spring-spawners when compared to (i) Atlantic spring-spawning herring populations from the Northern Atlantic (Table 1) or (ii) when compared to Baltic autumn-spawners (Figs 3c and 4c). To detect if introgressed regions are under selection in Baltic spring-spawning herring when compared to Atlantic spring-spawning herring, we revisited the pool-sequencing data from ^14^ and ^13^. In particular, we use genome-wide differentiation scans between North Atlantic spring-spawners and Baltic Sea spring- and autumn-spawners to determine if introgression show a strong signal of positive selection in the Baltic populations (Table 1). To further differentiate signatures of selection between Baltic Sea spring- and autumn-spawners we use Cross-population Extended Haplotype Homozygosity (xp-EHH) ^60^.

### Branch selection analysis with PAML

To test for lineage-specific positive selection in *SEC16B*, we performed branch model analyses using codeml implemented in PAML v4.9e ^61^. Coding sequences for the longest *SEC16B* isoform (ENSEMBL 109) were obtained for European sprat, rainbow trout (*Oncorhynchus mykiss*), and zebrafish (*Danio rerio*), along with two Atlantic herring individuals representing distinct haplotypes. The S4TM211 individual (Canada, Atlantic) represented the Atlantic spring-spawning haplotype, while the F1 Baltic Spring individual represented the Baltic spring-spawning haplotype.

Amino acid sequences were aligned using Clustal Omega ^62^ implemented in R through the ape package (v5.8-1) ^63^. Codon alignments were generated from the amino acid alignments using PAL2NAL ^64^ to preserve codon structure. A guide phylogeny was pruned from a published actinopterygian tree from the Fish Tree of Life project ^65^ to include only the four species analyzed (*C. harengus, S. sprattus, O. mykiss*, and *D. rerio*), and branch lengths were removed prior to use in PAML.

We applied branch models in codeml to estimate the ratio of nonsynonymous to synonymous substitution rates (ω = dN/dS) across lineages. The null model (M0) assumed a single ω across all branches. Alternative models allowed the ω ratio to vary on one or both herring branches (the Atlantic spring or Pacific herring allele lineages) while keeping all background branches constrained to a shared ω. Likelihood ratio tests (LRTs) were used to compare nested models (e.g., one-ratio vs. two- or three-ratio models), with test statistics (2ΔlnL) evaluated against a χ^2^ distribution with degrees of freedom corresponding to the difference in the number of parameters. We conducted three sets of tests: (1) with both herring branches designated as foreground, (2) with only the Atlantic spring allele branch as foreground, and (3) with only the Pacific herring allele branch as foreground. Log-likelihood values (lnL) were extracted from codeml output, and LRTs were computed in R.

### Annotation of a retin-specific THRB-isoform based on zebrafish annotation

The zebrafish THRB2 isoform is highly expressed in the retina and regulates the expression of long-wave opsins. THRB2 is distinguished from other THRB isoforms in zebrafish by the N-terminus encoded by a single conserved exon ^24^. We used reported sequences of exon 1 in the literature ^24,25^ to identify THRB2 as zebrafish isoform THRB-205 in ENSEMBL 114 (Accessed August 11 2025). We used the nucleotide sequence of exon 1 and blasted it to the Atlantic herring genome in ENSEMBL 109 (annotation used in this work) and 114 (most recent Atlantic herring annotation) using default settings. This allowed us to identify thrb-203 (ENSEMBL 109), and THRB-206 or THRB-207 (ENSEMBL 114) as the herring THRB2 isoforms. THRB-206 and THRB-207 differ in the presence/absence of one exon. As both of these isoforms overlap with the zebrafish retina-specific exon, both could potentially be expressed in the herring retina, though this requires further functional validation. In any case, THRB-203 (ENSEMBL 109) and THRB-206 (ENSEMBL 114) have the same genomic sequence and are the same isoform albeit with different names in the two annotations. Furthermore, the sequence of the first exon of THRB where our candidate missense mutation chr19:6359267 is placed does not differ between annotations. For further confirmation, we used MAFFT with default options (online tool 12-08-2025) to align the sequence of the first exons of THRB-205 from zebrafish and THRB-203 (ENSEMBL 109), THRB-206 and THRB-207 (ENSEMBL 114).

### Expression of opsin genes in correlation with THRB genotype

Eighteen herring samples were used to extract RNA for gene expression profiling. The tissue of interest was the retina; spring spawning herrings were caught in Hästskär, Sweden, during normal fishing season on May 8th and June 11th 2024 (Supplementary Table 1), and dissected at the Uppsala Biomedical Centre. Prior to gene expression analysis it was necessary to genotype the animals for position chr19:6359267 to screen for the CAA to CAC mutation in codon 40 ([ENSCHAT00020008343.1](https://www.ensembl.org/Clupea_harengus/Transcript/Sequence_cDNA?db=core;g=ENSCHAG00020003873;r=19:6346614-6359386;t=ENSCHAT00020008343;tl=YfwIU04udc48SMeS-11409446-1026869206), THRB-206), which corresponds to a change of Gln to His in the protein sequence. Genotyping was done by Sanger sequencing. DNA was extracted from fin clips with the DNeasy® 96 Blood&Tissue Kit (Cat. No. 69581, QIAGEN GmbH, Hilden, Germany) and three primer pairs were designed and tested to amplify the region surrounding the codon of interest. Primer pair forward TGCCCGACATGTATGAGATGC and reverse CGCTCATACCAGAGAAAGGTCC was used to generate a 402-bp PCR product with the KAPA HiFi PCR Kit (KK2102, Kapa Biosystems Pty Ltd, Cape Town, South Africa). AMPure XP beads (A63881, Beckman Coulter, Inc., Brea, USA) were used to purify the amplicons with a 1.1X ratio of beads to PCR volume. Samples were sent to Eurofins Genomics (Cologne, Germany) in a final volume of 17 µL, of which 15 µL were composed of 75 ng of purified PCR product and 2 µL of sequencing primer at a concentration of 10 µM. The software SnapGene® (Dotmatics, available at www.snapgene.com) was used to analyze the .ab1 and .seq files; the .fasta files were used for alignment to the Atlantic herring reference genome. From the May sampling batch nine homozygotes and six heterozygotes were selected to proceed for LSW1 and LSW2 expression, whereas three heterozygotes were picked from the June sampling batch.

RNA was extracted using a commercial kit (QIAGEN, 73404) following manufacturer’s instructions. Following RNA extraction, cDNA was synthesized using a cDNA synthesis kit (Thermo Fisher Scientific, 4368814) according to the manufacturer’s instructions. The synthesized cDNA was then used for subsequent real-time quantitative PCR (qPCR) analysis (Thermo Fisher Scientific, 4309155) on an Applied Biosystems QuantStudio 6 real-time PCR instrument. The PCR reaction program was set as follows: initial denaturation at 95°C for 10 min, followed by 40 cycles of denaturation at 95°C for 15 s, annealing at 60°C for 30 s, and extension at 72°C for 30 s. At the end of the reaction, a melting curve analysis was performed with the following settings: 95°C for 15 s, 60°C for 1 min, and 95°C for 15 s. The qPCR data were analyzed using the 2^-ΔΔCT^ method for relative quantification, with ACTIN and EF1A as internal controls. The CT values of the target genes LSW1 and LSW2 were compared with those of the internal control genes to calculate gene expression levels. We used a t-test to find significant differences in expression between genotypes. The primer sequence used were as follows: Detailed qPCR results in Supplementary Table 8.

### Correlation of THRB and RHO missense mutations

To explore the genetic correlation between *THRB* and *RHO* missense mutations, genotype data of 150 Baltic herring populations were accessed from European Variant Archive (EVA) accession PRJEB96176 (Goodall et al. 2025, *in review at PNAS*^*28*^). Allele frequencies for both reference and alternate alleles of *THRB* and *RHO* were calculated using PLINK (v.1.9, ^66^, as per Goodall et al. (2025)^28^. Pairwise linear models between *THRB* (H40Q allele, chr 19:6359267) and *RHO* F261Y (chr 4:11217516) and T213I (4:11217660) alleles were calculated in *R* alongside summary statistics, including adjusted R^2^ and *P* value using a linear regression with the function *lm()* in R. For all comparisons, spawning ecotypes (spring-, summer-, and autumn-spawners) were defined relative to time of capture (Goodall et al. 2025, *in review at PNAS*^*28*^) where individuals captured before July were denoted as spring, during July as summer, and after July as autumn.

## Acknowledgements

The authors thank members of the Leif Andersson group for valuable comments and discussions on the preliminary results. In particular, we thank Ryan Daniels for sample collection and Giulia Maestri for assistance with experimental work. This study was supported by the Knut and Alice Wallenberg Foundation (KAW 2023.0160) and the Swedish Research Council (Vetenskapsrådet; 2017-02907). Computational infrastructure was provided by the National Academic Infrastructure for Supercomputing in Sweden (NAISS), partially funded by the Swedish Research Council through grant agreement no. 2022-06725. M.S.F. was supported by a Marie Skłodowska-Curie European Postdoctoral Fellowship (Project 101063864, INVERT2ADAPT) awarded by the European Research Executive Agency, and by a Science for Life Laboratory Fellowship.

## Data availability

All genomic data used in this manuscript have been previously published and are deposited in NCBI under the following IDs: SAMN41434462, PRJNA930418, PRJNA642736, PRJNA1023520, PRJNA338612, SRA057909, and PRJEB32358 (see Supplementary Table 1). Sequence data for the *THRB* locus for 18 individuals will be available on Figshare. The original code used for the analyses will be made available on GitHub (https://github.com/MafaldaSFerreira/Baltic_herr_introgression) and permanently archived in Zenodo. Intermediate files used in the analyses will also be available on Github and Figshare.

