## Supplementary Figures and Tables for "Adaptive introgression from Pacific herring to Atlantic herring in the brackish Baltic Sea"

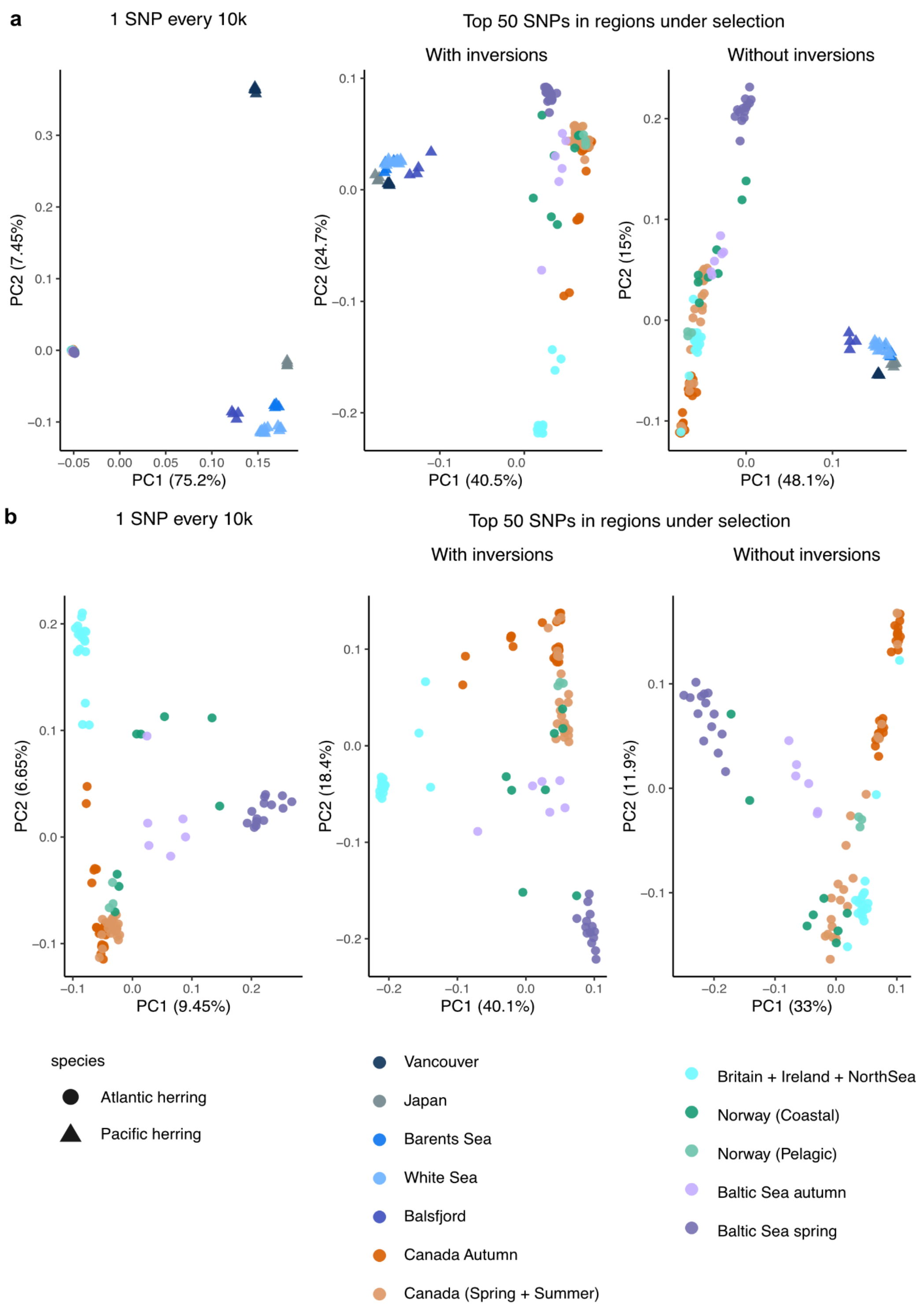


**Supplementary Fig. 1 - Population structure in Atlantic and Pacific herring.** Principal component analysis of single nucleotide variation data for **a)** 125 Pacific herring and Atlantic herring individuals or **b)** only Atlantic herring individuals.

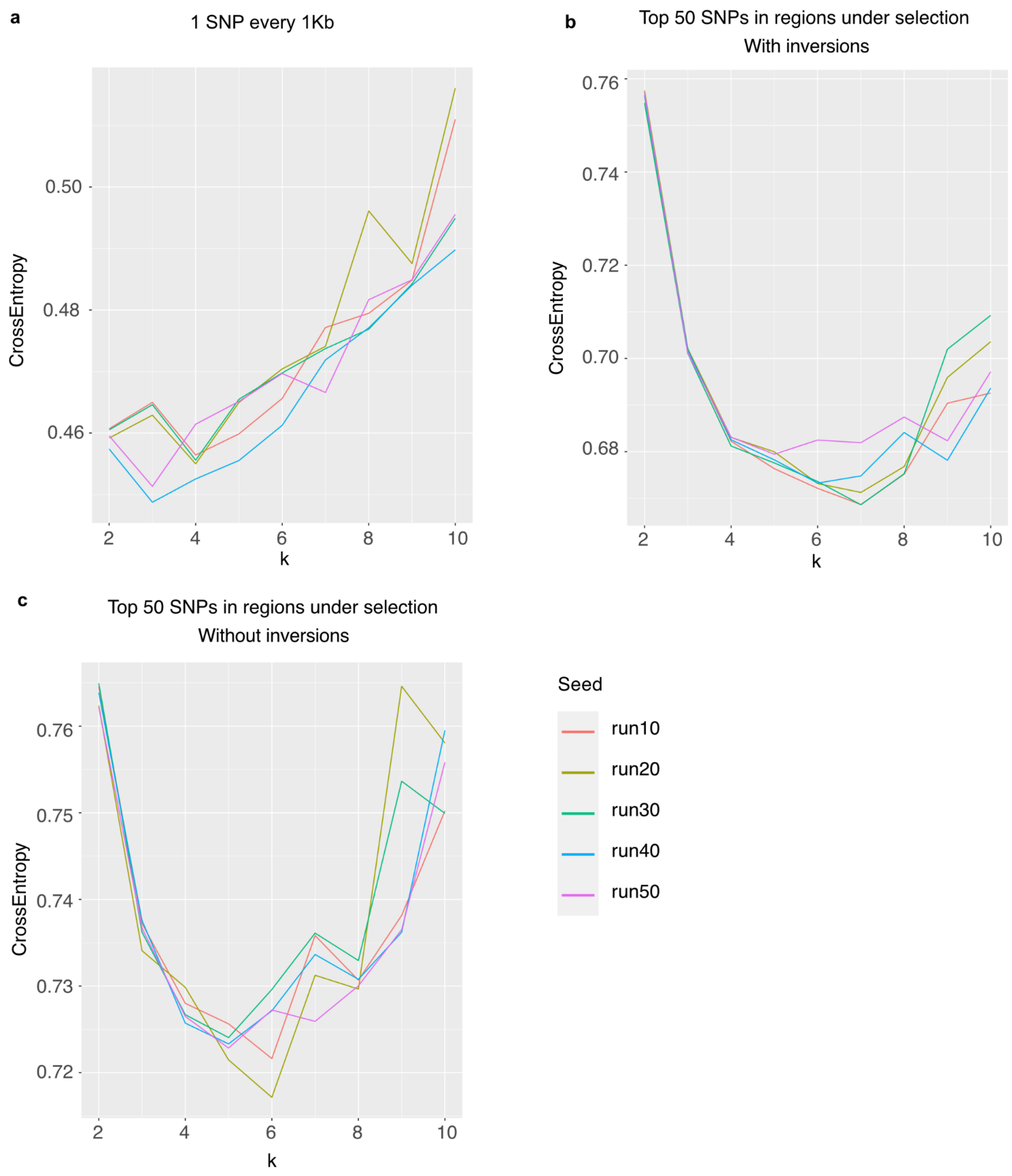


**Supplementary Fig. 2 -** Cross validation error analysis for ancestry analysis performed with sNMF with genetic data for 125 Atlantic and Pacific herring individuals. The analysis was run with three different sets of genetic variants: **a** 67,566 SNPs spaced at least 1 Kb apart; **b** 15,231 SNPs within the top regions under selection,including inversion regions; and **c** 11,492 SNPs within regions under selection, excluding inversion regions.

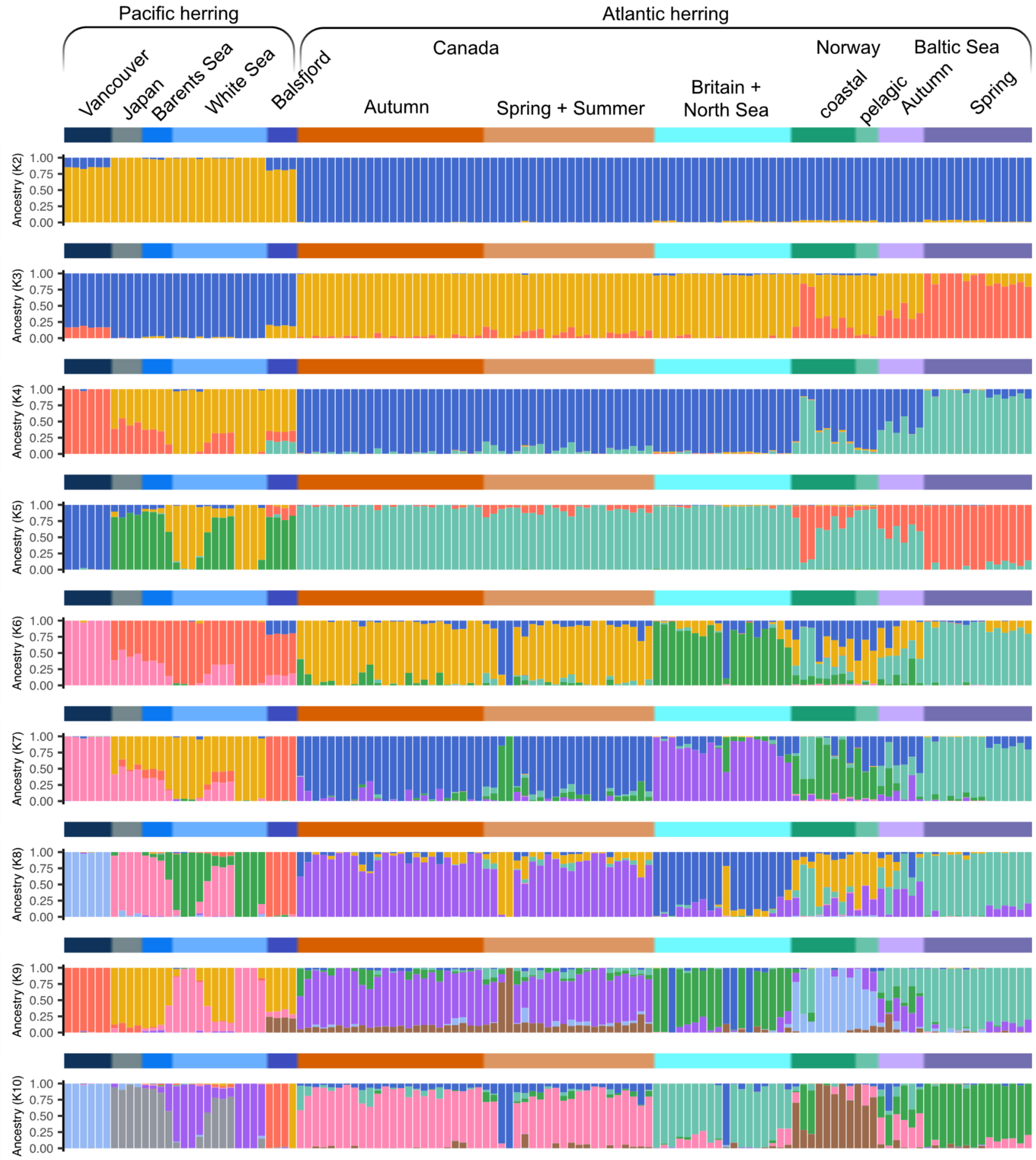


**Supplementary Fig. 3 -** Ancestry analysis performed with sNMF and genetic data for 125 Atlantic and Pacific herring individuals. The analysis was run with SNPs distancing at least 1 KB from each other.

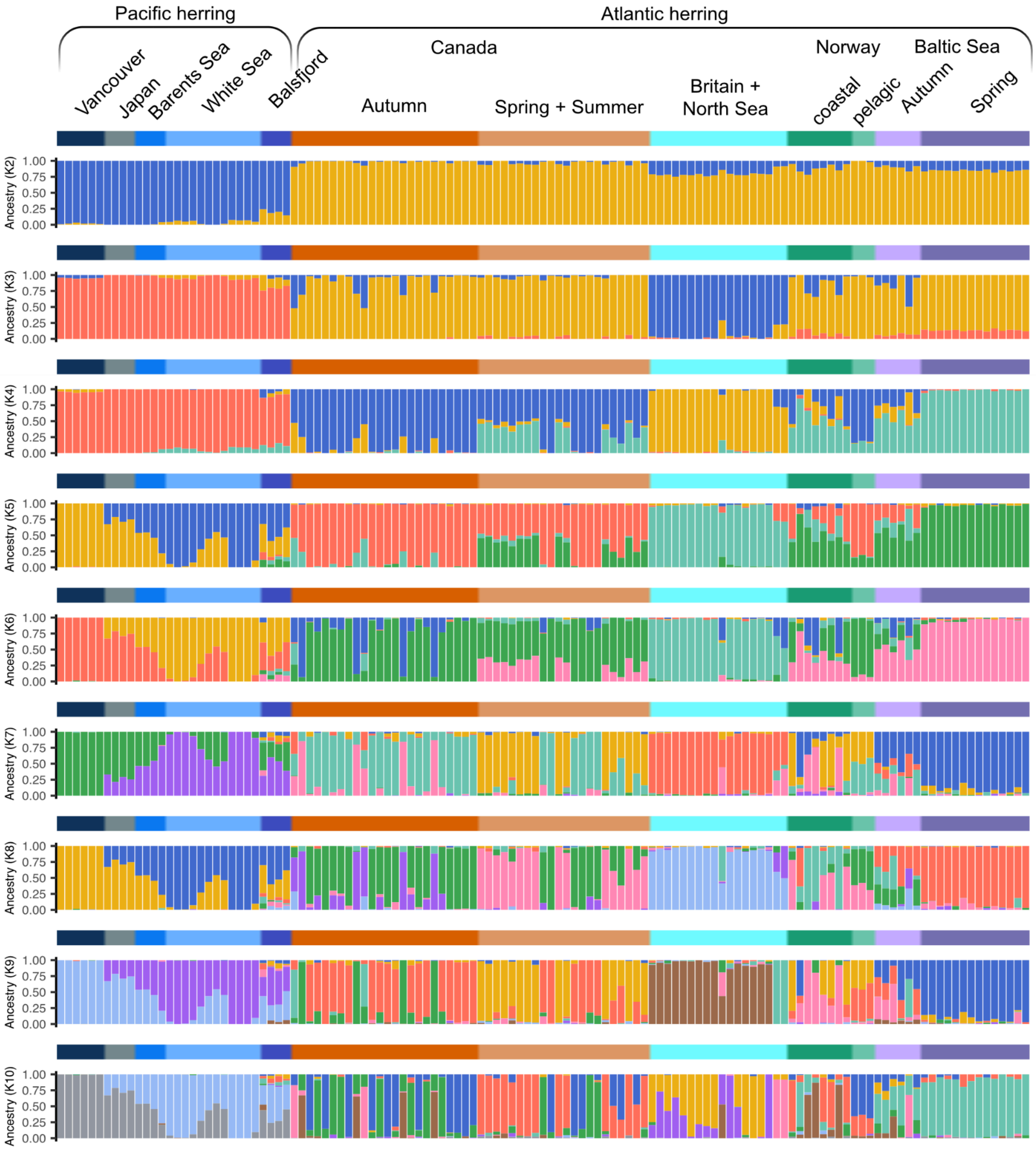


**Supplementary Fig. 4 -** Ancestry analysis performed with sNMF and genetic data for 125 Atlantic and Pacific herring individuals. The analysis was run with SNPs located in regions under selection including inversions regions.

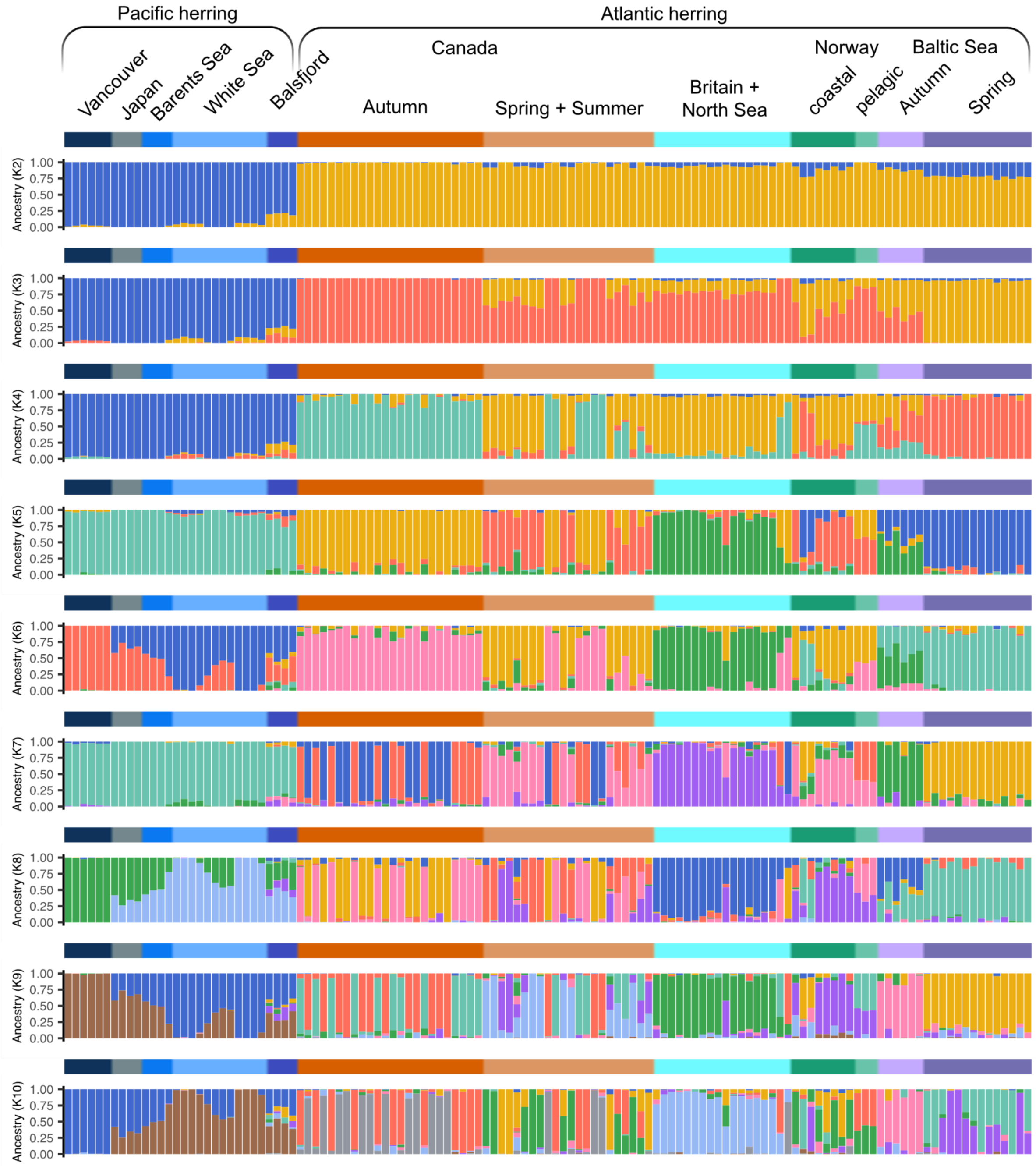


**Supplementary Fig. 5 -** Ancestry analysis performed with sNMF and genetic data for 125 Atlantic and Pacific herring individuals. The analysis was run with SNPs located in regions under selection excluding inversions regions.

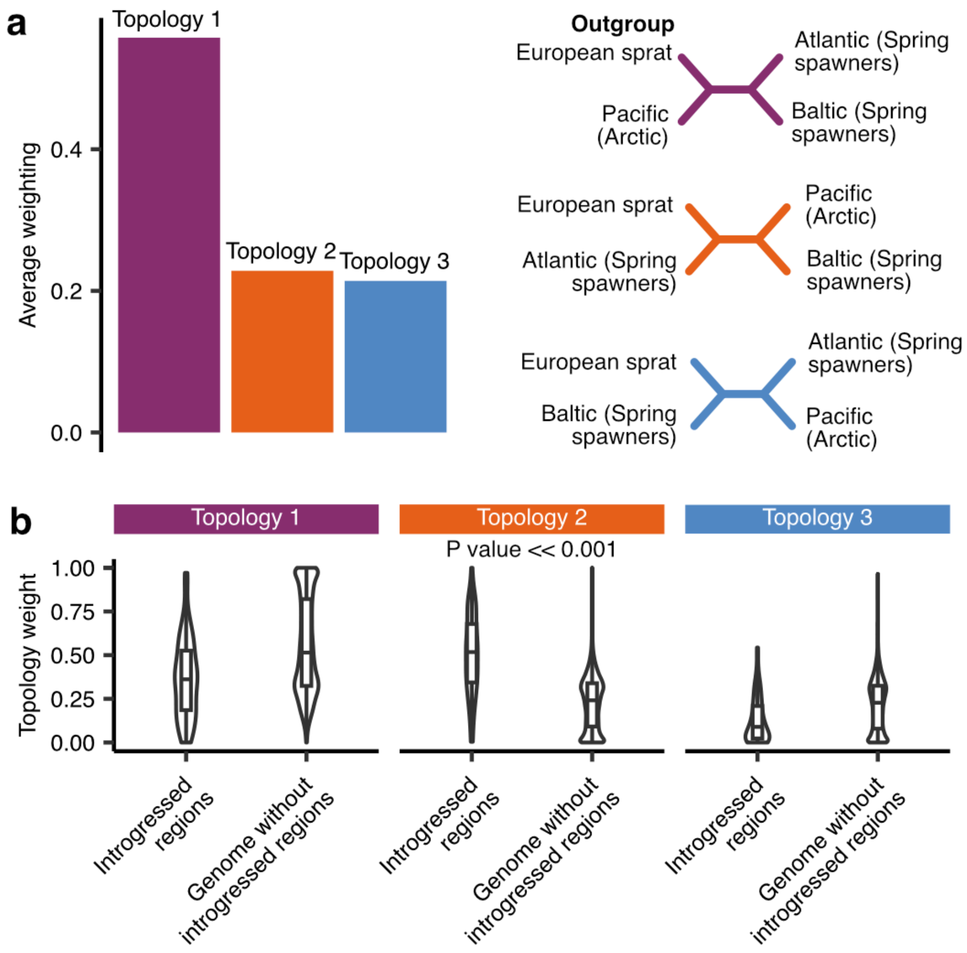


**Supplementary Fig. 6 -** Twisst summary results. Average support for topology 2 in the background and in 8-fold recurring Atlantic HSRs, respectively.

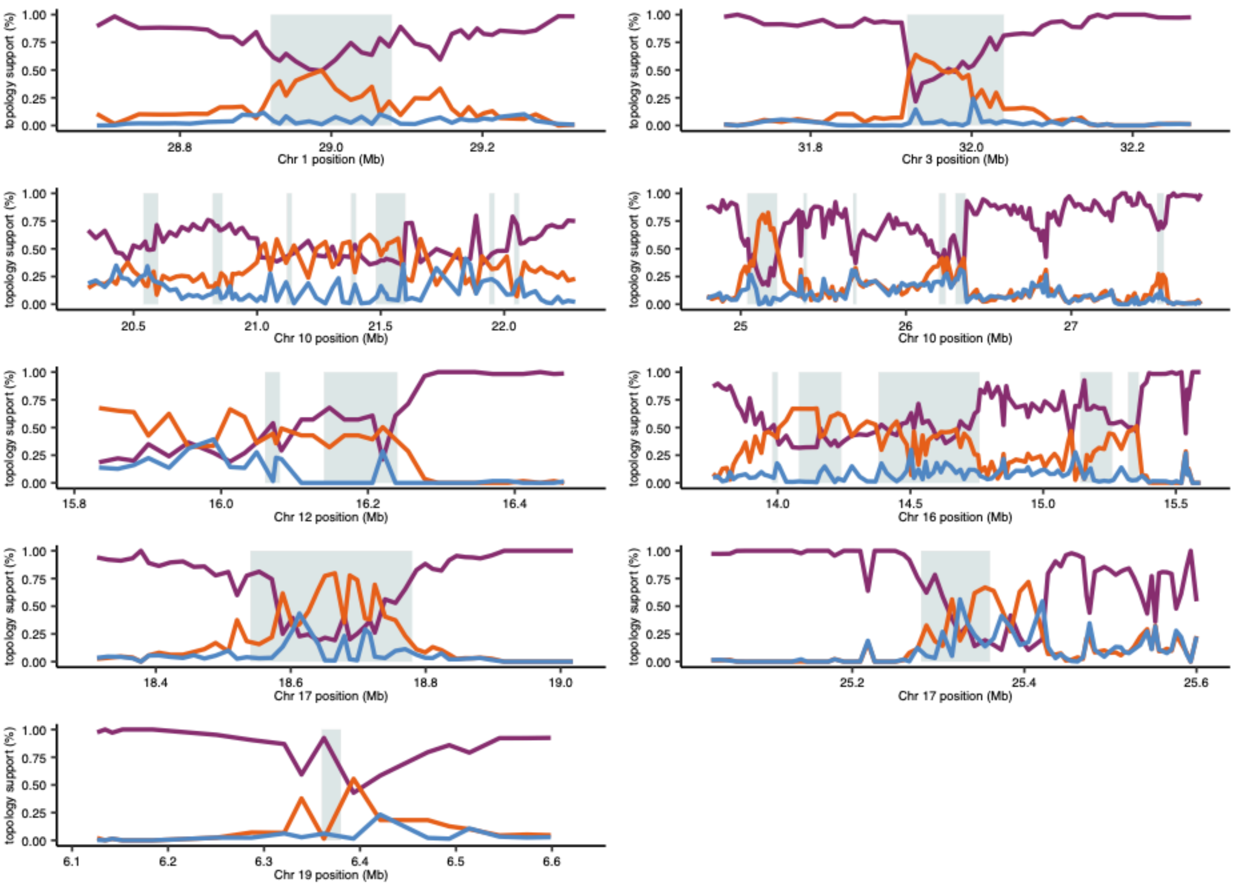


**Supplementary Fig. 7 -** Topology analysis with Twisst using Pacific herring individuals from Vancouver as an outgroup. Introgression regions identified by a genome-wide scan (Fig. 2) are shown, with significant introgressed regions highlighted in light blue. The purple, orange and blue lines represent the weight of each alternative topology in sliding windows of 100 SNPs.

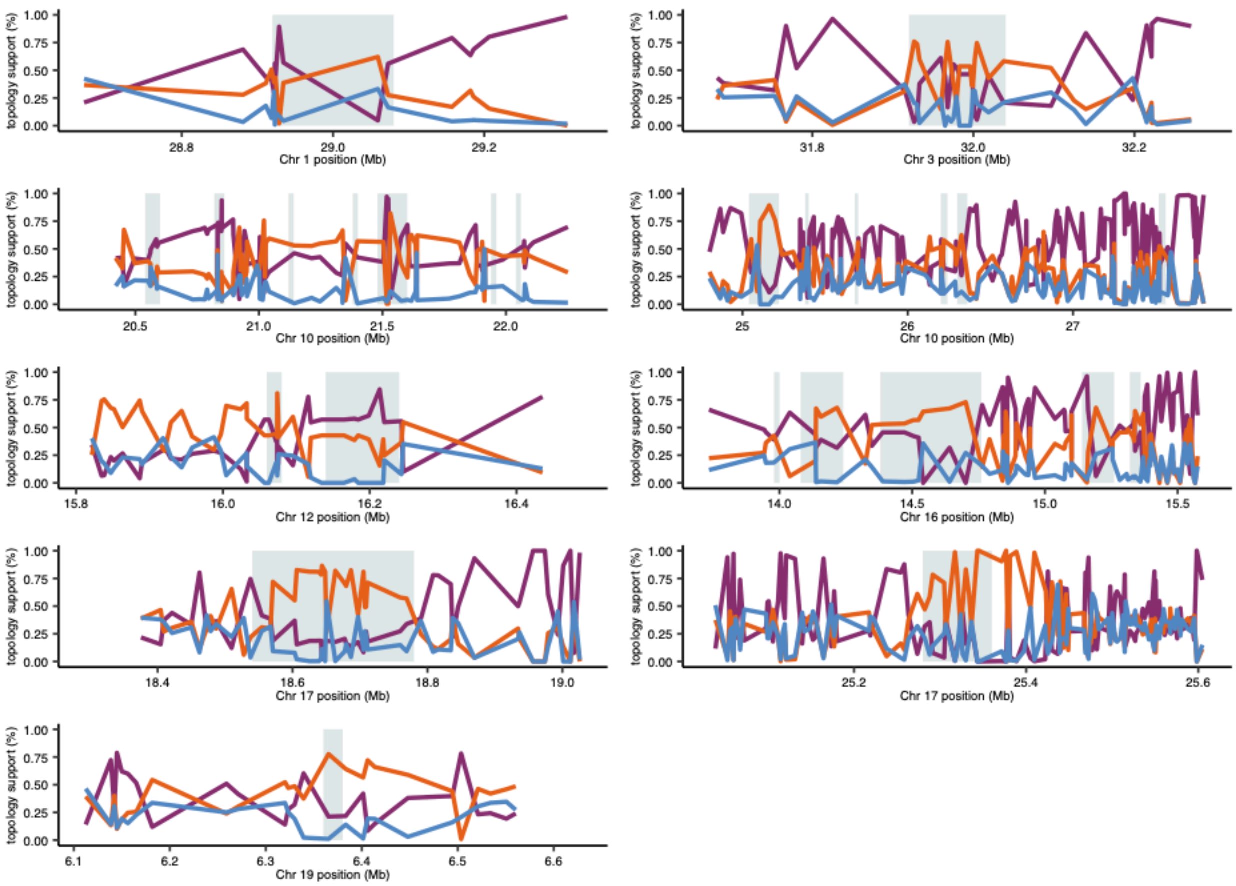


**Supplementary Fig. 8 -** Topology analysis with Twisst using European sprat as an outgroup. Introgression regions identified by a genome-wide scan (Fig. 2) are shown, with significant introgressed regions highlighted in light blue. The purple, orange and blue lines represent the weight of each alternative topology in sliding windows of 100 SNPs.

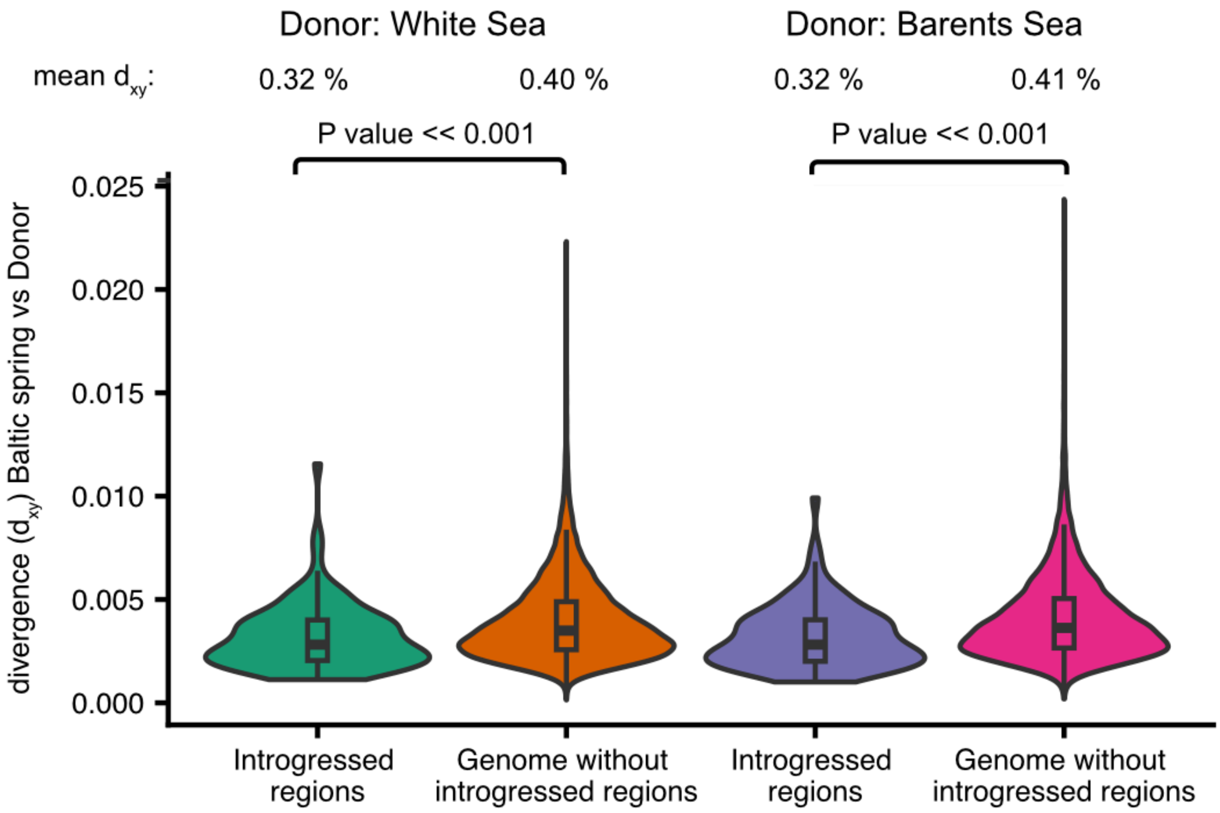


**Supplementary Fig. 9 -** Absolute divergence ($d_{xy}$) in introgression and non-introgressed regions of the genome between Baltic spring-spawning herring individuals and the two putative Arctic Pacific herring donor populations, White Sea and Barents Sea. T-tests were performed to test the significant difference between $d_{xy}$ distribution between introgressed and non-introgressed regions.

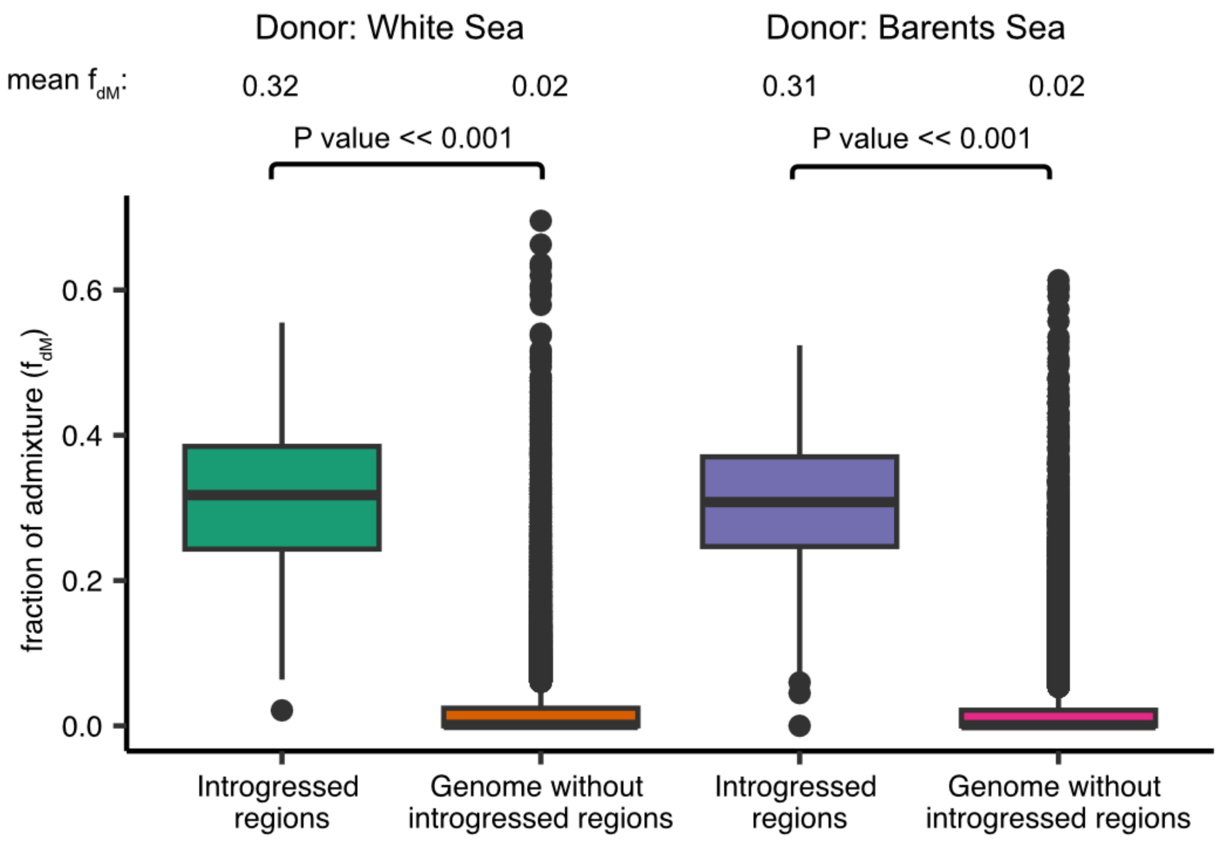


**Supplementary Fig. 10 -** Proportion of introgression ($f_{dM}$) in introgressed and non-introgressed regions of the genome between Baltic spring-spawning herring individuals and the two putative Arctic Pacific herring donor populations, White Sea and Barents Sea. T-tests were performed to test the significant difference between $f_{dM}$ distribution between introgressed and non-introgressed regions.

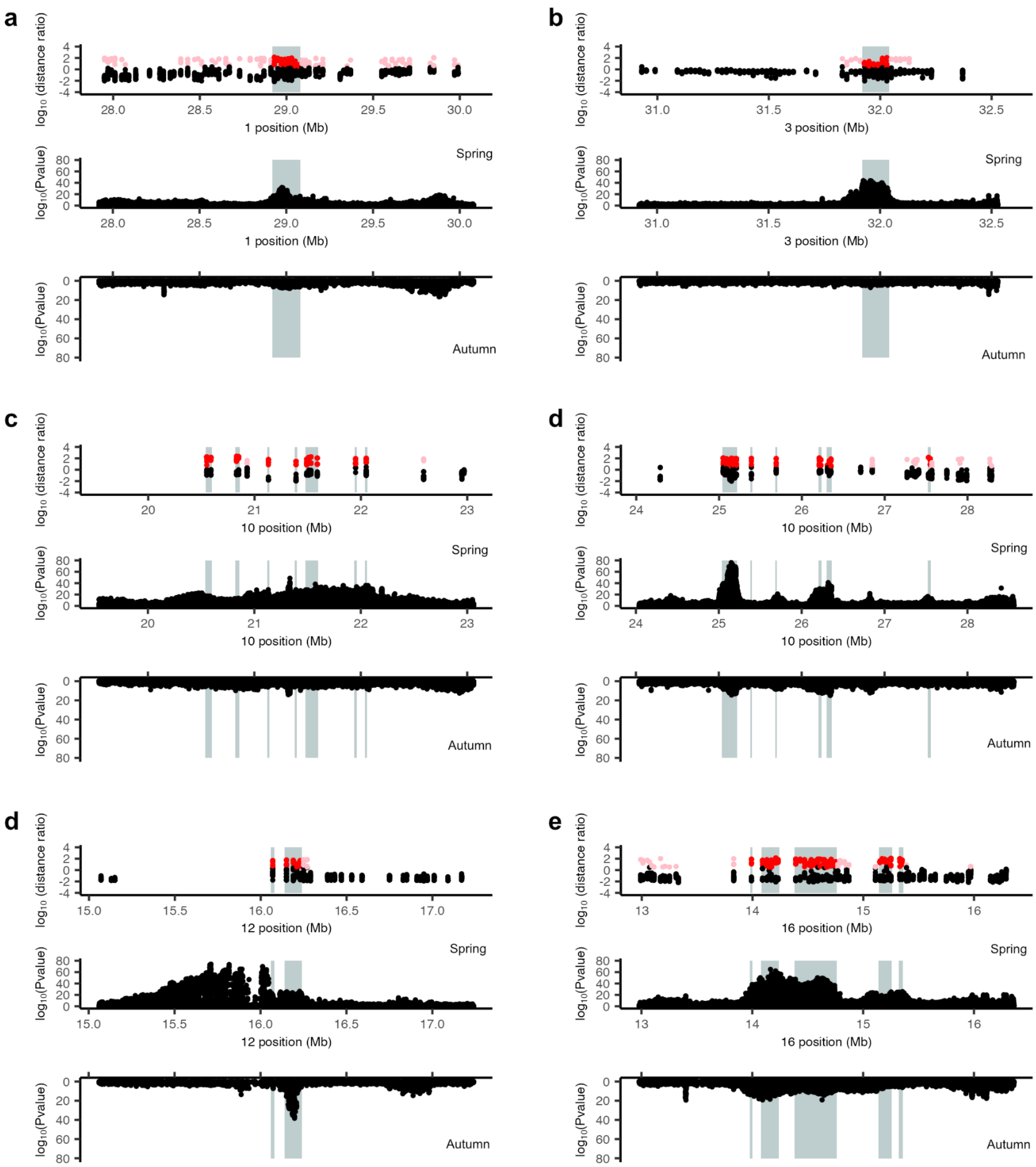


Supplementary Fig. 11 (continues in the next page)

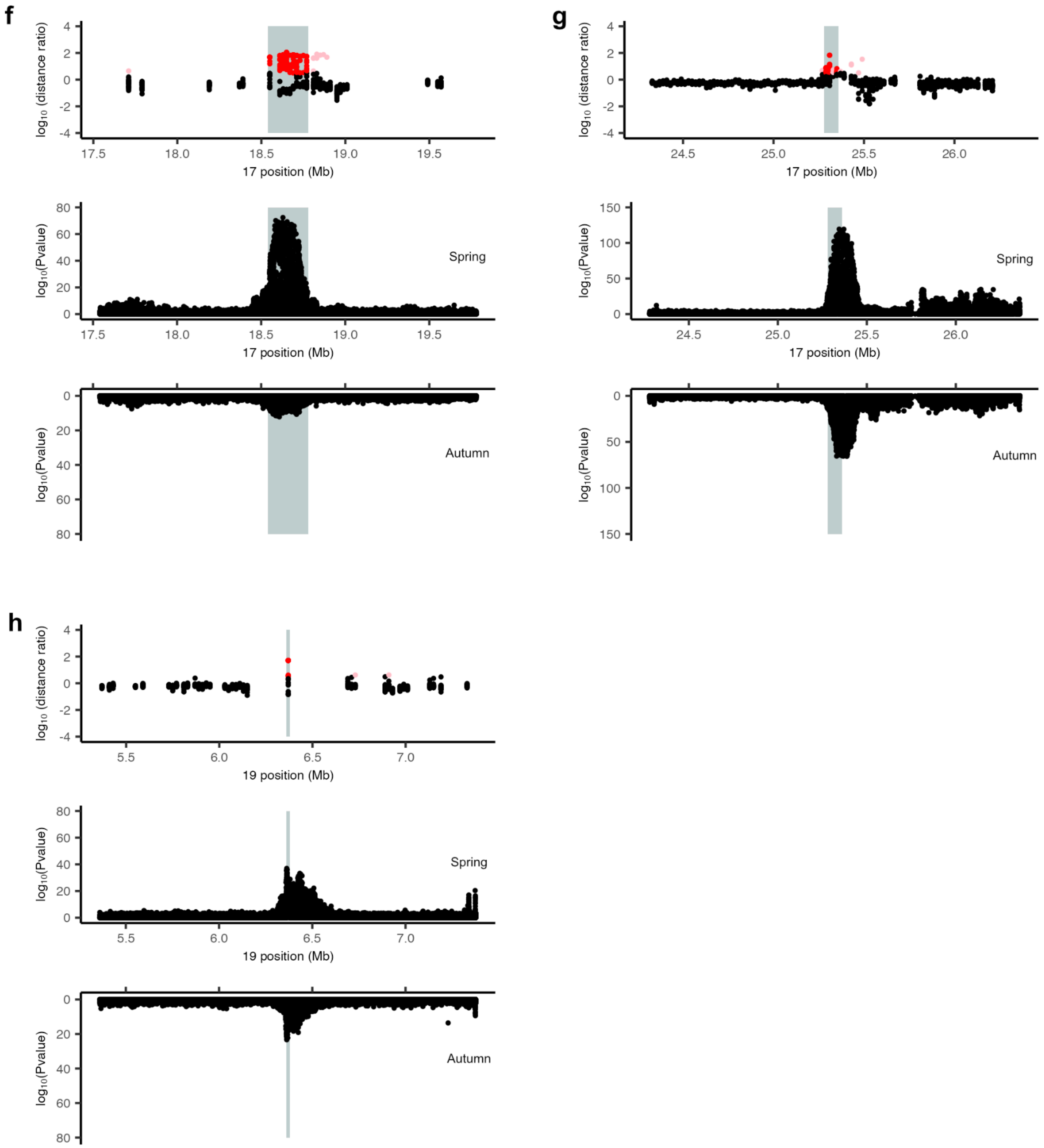


**Supplementary Fig. 11 -** Overlap between introgression scan results and allele frequency differentiation scan results. **a-h** For each region of introgression, we show the results of the introgression scan testing for introgression between Baltic spring-spawning herring and Arctic pacific herring (top panel), the p-value for the test of allele frequency differentiation between Atlantic and Baltic spring-spawning herring (middle panel) or Atlantic and Baltic autumn-spawning herring (bottom panel). Introgression regions are highlighted in gray boxes. In the introgression scan, dots represent 20 kb windows. Pink and red dots represent significantly introgressed windows based on distance or distance and coverage (seven haplotypes introgressed), respectively.

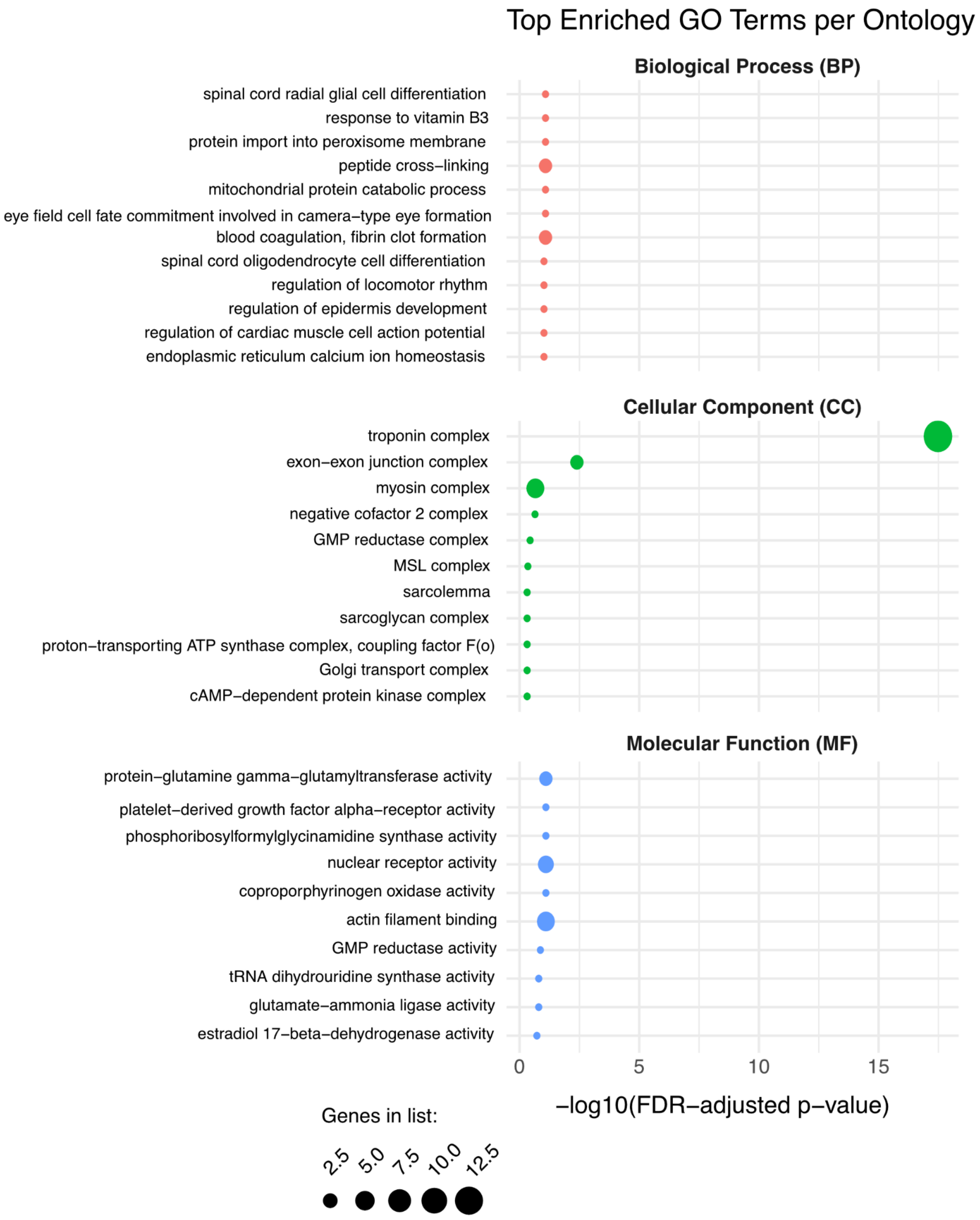


**Supplementary Fig. 12-** Gene Ontology enrichment analysis for 166 genes overlapping with introgression regions. The Top 10 terms are presented for each GO Term Category: Biological Process (BP), Cellular Component (CC) and Molecular Function (MF). We used topGO (2.58.0) with the weighted01 algorithm and Fisher’s exact test to determine significance.

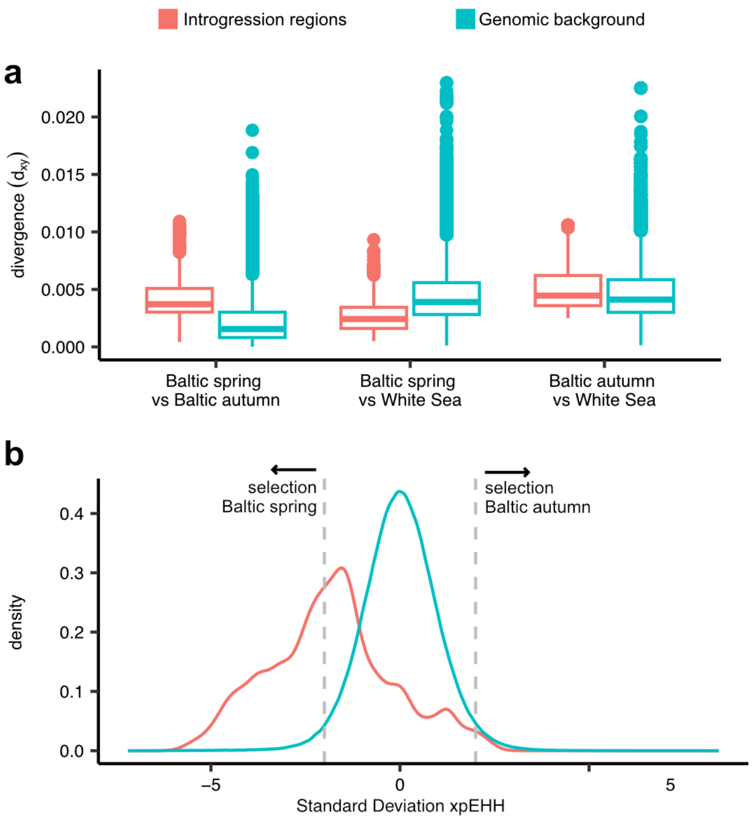


**Supplementary Fig. 13 -** Divergence to the Arctic Pacific herring (here White Sea chosen as representative) and selection in introgressed regions (in salmon color) and genomic background (in blue) in Baltic Sea spring- and autumn-spawners. **a** Divergence between populations as indicated on the x-axis for introgression regions compared with genomic background; **b** Standardized Cross population Extended Haplotype Homozygosity (a measure of positive selection) in introgressed regions and genomic background. Values of xpEHH below -2 indicate positive selection in Baltic spring-spawners and above 2 indicate selection in Baltic autumn-spawners.

**a**

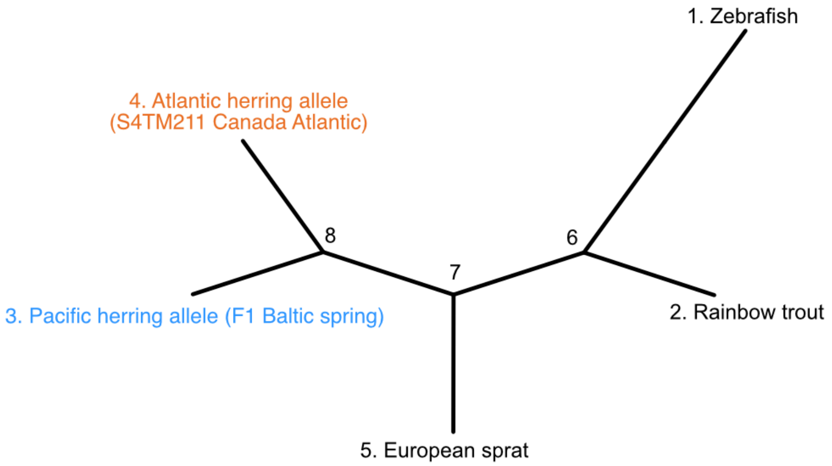


**b**

| **Hypothesis** | **lnL** |
|---|---|
| H1: w0 (dN/dS is equal for all branches) | -9299.0 |
| H2: w1 for Atlantic herring allele = AS and for other branches, 2 w | -9295.8 |
| H3: w2 for Pacific herring allele = BS and for other branches, 2 w | -9298.6 |
| H4: w1, w2 and 1 dN/dS for other branches, 3 w | -9295.1 |

| **Likelihood Ratio Test** | **P-value** |
|---|---|
| w0=w1=w2 vs w0=w1≠w2 (All evolve at same w) vs (selection in BS) | 0.349 |
| w0=w1=w2 vs w0=w2≠w1 (All evolve at same w) vs (selection in AS) | 0.011 |
| w0=w1=w2 vs w0≠w1≠w2 (All evolve at same w) vs (selection in BS and AS) | 0.020 |
| w0=w1≠w2 vs w0≠w1≠w2 (selection in BS) vs (selection in BS and AS) | 0.008 |
| w0=w2≠w1 vs w0≠w1≠w2 (selection in AS) vs (selection in BS and AS) | 0.245 |

**c** Hypothesis 4: Three rates of selection

| **Branch** | t | N | S | dN/dS | dN | dS | N*dN | S*dS |
|---|---|---|---|---|---|---|---|---|
| 6..1 | 1.19 | 1893.1 | 653.9 | 0.2245 | 0.2102 | 0.9362 | 397.9 | 612.2 |
| 6..2 | 1.512 | 1893.1 | 653.9 | 0.2245 | 0.2671 | 1.1895 | 505.6 | 777.8 |
| 6..7 | 1.059 | 1893.1 | 653.9 | 0.2245 | 0.1871 | 0.8335 | 354.3 | 545.1 |
| 7..8 | 0.044 | 1893.1 | 653.9 | 0.2245 | 0.0078 | 0.0349 | 14.9 | 22.8 |
| 8..4 | 0.011 | 1893.1 | 653.9 | 1.7366 | 0.0042 | 0.0024 | 7.9 | 1.6 |
| 8..3 | 0.002 | 1893.1 | 653.9 | 0.0001 | 0 | 0.0023 | 0 | 1.5 |
| 7..5 | 0.094 | 1893.1 | 653.9 | 0.2245 | 0.0167 | 0.0742 | 31.5 | 48.5 |

**Supplementary Fig. 14** - Branch model analysis from codeml testing for different hypotheses of evolution for *SEC16B*. **a** Phylogenetic tree used to test for branch models in PAML. Tips and nodes are labeled with numbers that correspond to branches in panel (c). **b** Likelihood for each hypothesis tested with codeml, assuming different ω (ratio of nonsynonymous to synonymous substitution rates; or dN/dS) ratios. We test a one-ratio model (H1), two two-ratio models (H2 and H3), and a three-ratio model (H4). We used a likelihood ratio test to determine which hypothesis fits best. **c** Output for the three-ratio model (H4), which allows both Pacific herring and Atlantic herring alleles to have a different dN/dS than the rest of the tree.

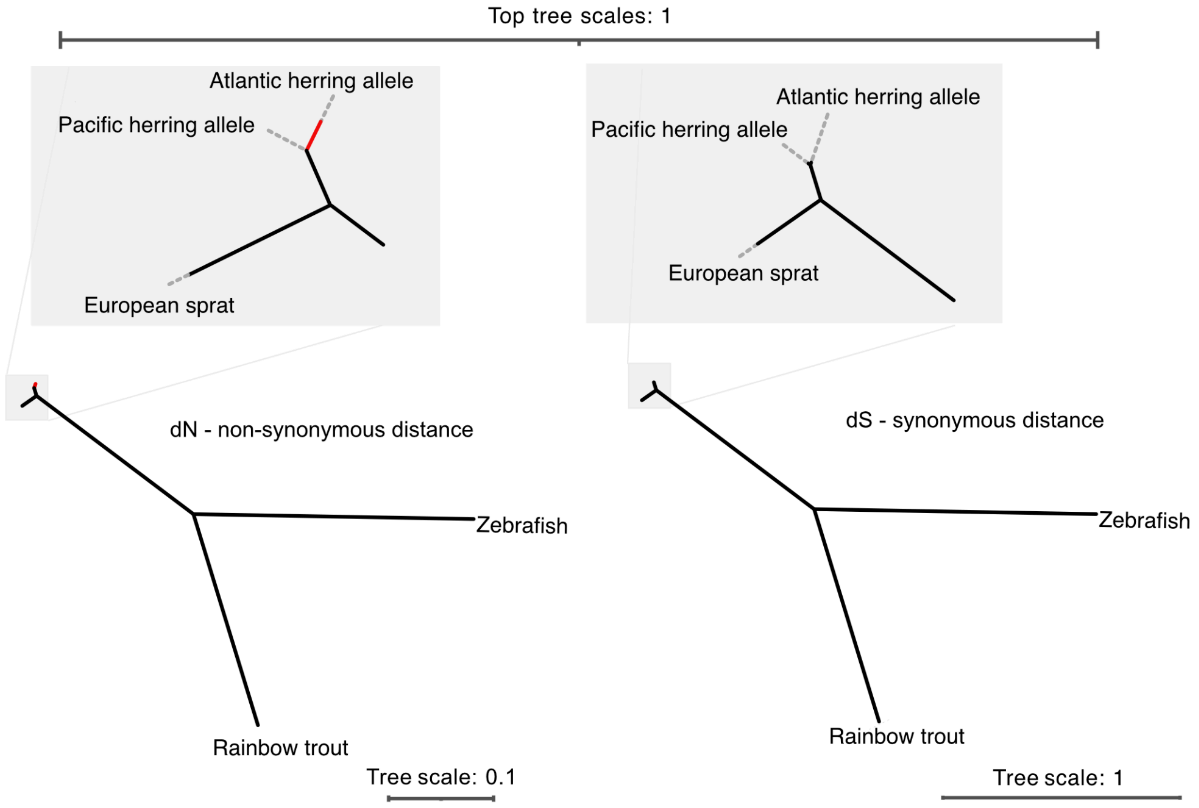


**Supplementary Fig. 15 -** dN and dS phylogenetic trees for *SEC16B*. Insets provide zoom-ins into the herring and European sprat clade, evidencing the difference in branch length between the Pacific herring allele and Atlantic herring allele of *SEC16B*. Branch lengths are provided in Supplementary Fig. 14c.

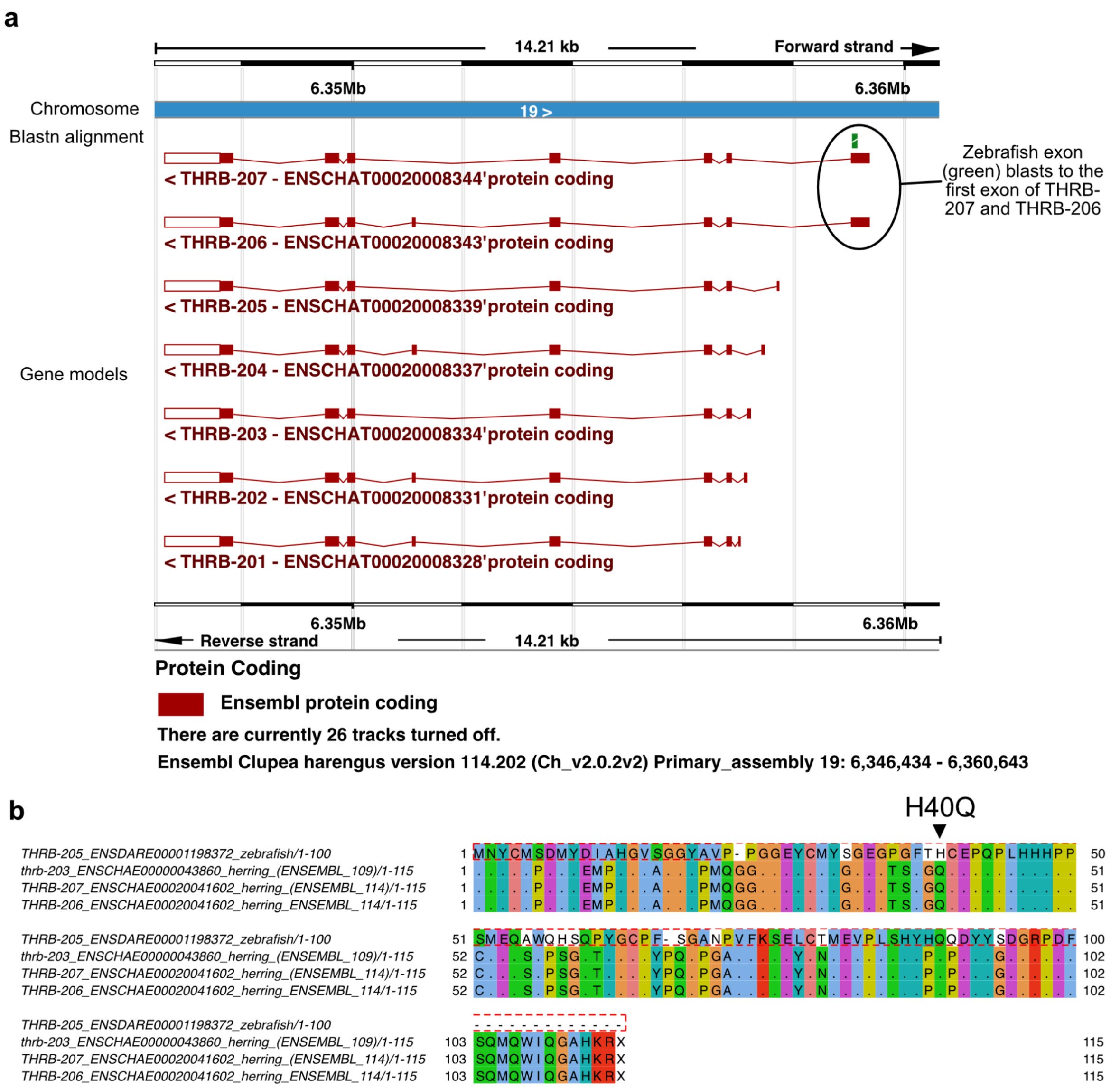


**Supplementary Fig. 16 -** Annotation of the retina specific THRB isoform in Atlantic herring. **a** Blastn alignment of the first exon of the retina specific THRB isoform from zebrafish to the Atlantic herring genome (ENSEMBL 114). The zebrafish exon (green) blasts to the first exon of the THRB-207 and THRB-206 isoforms in herring, identifying these two as the putative retina specific THRB isoforms in herring. **b** Amino-acid alignment between the first exon of the retina specific isoform of zebrafish (THRB-205), and the first exons of the retina specific isoforms in Atlantic herring in the ENSEMBL 109 annotation (used to interpret functions of SNPs and genes in this work) and ENSEMBL 114 annotation (the most recent Atlantic herring annotation). The location of the H40Q amino-acid mutation is highlighted. The Atlantic herring reference genome has the Baltic spring genotype.

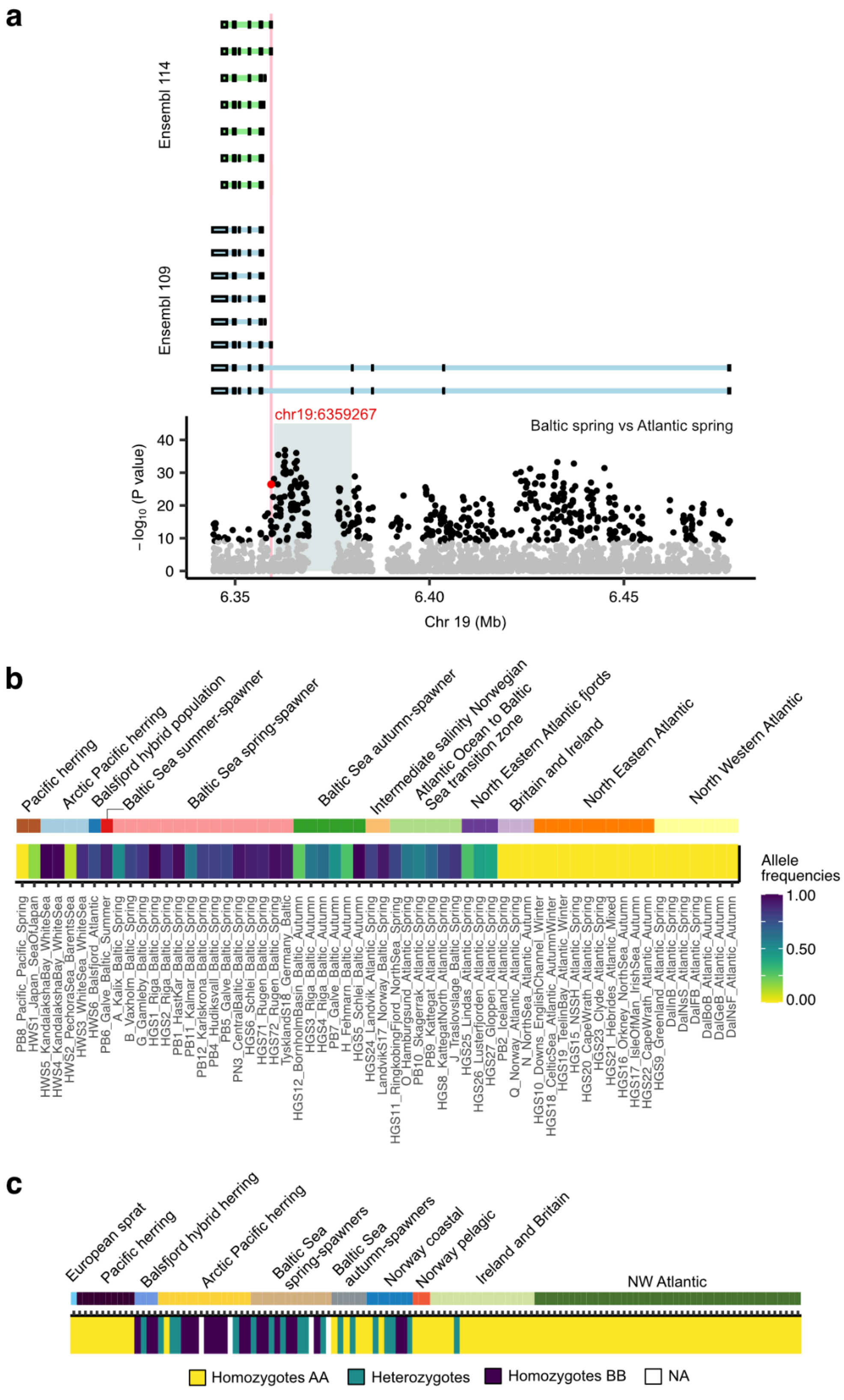


**Supplementary Fig. 17 (legend in the next page)**

**Supplementary Fig. 17 -** The H40Q *THRB* missense mutation in Atlantic herring. **a** Location of the H40Q missense mutation (red dot) relative to the ENSEMBl 109 and ENSEMBL 114 Atlantic herring annotation and the introgression region that overlaps with *THRB* (gray box). In both annotations, the missense mutation is located on the first exon of one or two isoforms that is not shared with the remaining isoforms. In zebrafish, the retina-specific THRB isoform (THRB2) has the same gene model, where the first exon is unique to the isoform expressed in the retina. **b** Allele frequency of the *THRB* H40Q missense mutation across Atlantic herring populations (using pool-seq data from Han et al 2020, eLife). Most populations in the Atlantic are fixed for one allele. Allele frequency variation only exists for populations distributed in the transition zone Atlantic Ocean and Baltic Sea or in the Arctic. **c** Genotype heatmap for the *THRB* H40Q mutation. The derived mutation is shared between Arctic Pacific herring and Baltic Sea spring-spawning herring, while the ancestral mutation is present in European sprat, Pacific Ocean Pacific herring and most Atlantic herring individuals.

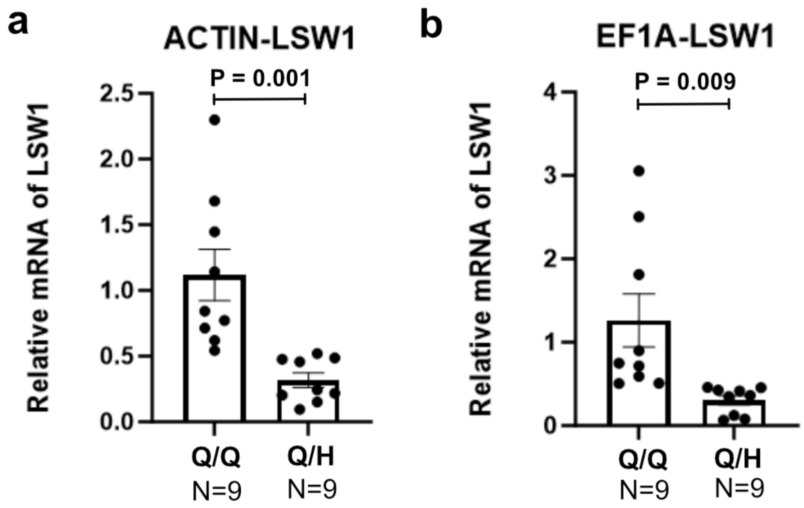


**Supplementary Fig. 18 -** Quantitative-PCR results for gene expression of *LSW1* in homozygotes and heterozygotes for the H40Q THRB missense mutation using **(a)** *ACTIN* or **(b)** *EF1A* genes as reference housekeeping genes. Significance differences between genotypes were tested using a t-test.

**Supplementary Table 1 -** Sample metadata for the sequencing data used in the present study.

**Supplementary Table 2 -** Percentage of introgression in Atlantic herring populations, considering at least seven introgressed haplotypes and merging the ones within 50 kb from each other.

| Target population | R File | Parent 1 | Parent 2 | Percentage |
|---|---|---|---|---|
| Baltic Sea spring-spawners | Scan1 v01 | NSSH Norway + Canada Atlantic Spring | White Sea + Pechora | 0.29 % |
| Baltic Sea autumn-spawners | scan2_v01_baltic_alt_ref_summary_cov7_gr_min50kb | NSSH Norway + Canada Atlantic Spring | White Sea + Pechora | 0.01 % |
| Britain and Ireland | scan3_v01_UK_alt_ref_summary_cov7_gr_min50kb | NSSH Norway + Canada Atlantic Spring | White Sea + Pechora | 0.003 % |
| Atlantic Ocean autumn-spawners (Canada) | scan4_v01_AtlAut_alt_ref_summary_cov7_gr_min50kb | NSSH Norway + Canada Atlantic Spring | White Sea + Pechora | 0.00 % |
| Atlantic Ocean spring-spawners (Norway + Canada) | scan7_v01_AtlSpring_v_subarctic_alt_ref_summary_cov7_gr_min50kb | Canada Atlantic Autumn | White Sea + Pechora | 0.06 % |
| North Sea | scan10_v01_NorthSea_alt_ref_summary_cov7_gr_min50kb | NSSH Norway + Canada Atlantic Spring | White Sea + Pechora | 0.00 % |

**Supplementary Table 3 -** Nine introgressed regions in Baltic spring-spawning herring. For each introgressed window, we indicate the chromosome, start and end coordinate and width. We also indicate the position of the top differentiated SNP between Atlantic and Baltic within spawning groups (spring or autumn) from Han et al. (2020).

| Chromosome | Start (Mb) | End (Mb) | Width (Kb) | Top SNP position in spring | Top SNP position in autumn |
|---|---|---|---|---|---|
| 1 | 28.92 | 29.08 | 160 | 28,975,098 | 29,017,606 |
| 3 | 31.92 | 32.04 | 120 | 31,924,585 | 31,957,191 |
| 10 | 20.54 | 20.6 | 60 | 20,581,916 | 20,595,972 |
| | 20.82 | 20.86 | 40 | 20,850,721 | 20,824,255 |
| | 21.12 | 21.14 | 20 | 21,130,733 | 21,128,136 |
| | 21.38 | 21.4 | 20 | 21,390,557 | 21,386,784 |
| | 21.48 | 21.6 | 120 | 21,572,857 | 21,553,571 |
| | 21.94 | 21.96 | 20 | 21,959,848 | 21,955,566 |
| | 22.04 | 22.06 | 20 | 22,046,953 | 22,059,214 |
| 10 | 25.04 | 25.22 | 180 | 25,150,137 | 25,153,331 |
| | 25.38 | 25.4 | 20 | 25,382,814 | 25,390,702 |
| | 25.68 | 25.7 | 20 | 25,699,987 | 25,686,760 |
| | 26.20 | 26.24 | 40 | 26,217,604 | 26,201,033 |
| | 26.30 | 26.36 | 60 | 26,319,497 | 26,344,098 |
| | 27.52 | 27.56 | 40 | 27,521,243 | 27,527,212 |
| 12 | 16.06 | 16.08 | 20 | 16,065,802 | 16,073,917 |
| | 16.14 | 16.24 | 100 | 16,227,402 | 16,197,898 |
| 16 | 13.98 | 14 | 20 | 13,999,471 | 13,980,589 |
| | 14.08 | 14.24 | 160 | 14,164,286 | 14,092,534 |
| | 14.38 | 14.76 | 380 | 14,588,897 | 14,629,571 |
| | 15.14 | 15.26 | 120 | 15,252,445 | 15,174,789 |
| | 15.32 | 15.36 | 40 | 15,355,296 | 15,354,567 |
| 17 | 18.54 | 18.78 | 240 | 18,629,585 | 18,608,403 |
| 17 | 25.28 | 25.36 | 80 | 25,344,618 | 25,330,413 |
| 19 | 6.36 | 6.38 | 20 | 6,362,829 | 6,362,840 |

**Supplementary Table 4 -** Gene content of introgressed regions and 20 kb surrounding area.

| Chromosome | Start | End | ENSEMBL Gene ID | External gene name | Description |
|---|---|---|---|---|---|
| 1 | 28925984 | 28930751 | ENSCHAG00000012097 | ATP5MC1 | ATP synthase membrane subunit c locus 1 |
| 1 | 28937816 | 28948986 | ENSCHAG00000012169 | UBE2Z | ubiquitin conjugating enzyme E2 Z |
| 1 | 28964896 | 29009013 | ENSCHAG00000012263 | IGF2BP1 | insulin-like growth factor 2 mRNA binding protein 1 |
| 1 | 29017833 | 29024009 | ENSCHAG00000012653 | LOC105912842 | uncharacterized LOC105912842 |
| 1 | 29021873 | 29022007 | ENSCHAG00000012661 | snoRNA | |
| 1 | 29027219 | 29035348 | ENSCHAG00000012664 | NR1D1 | nuclear receptor subfamily 1 group D member 1 |
| 1 | 29040035 | 29047401 | ENSCHAG00000012702 | MSL1A | MSL complex subunit 1a |
| 1 | 29050591 | 29064623 | ENSCHAG00000012759 | CASC3 | casc3 exon junction complex subunit |
| 1 | 29076864 | 29115910 | ENSCHAG00000012776 | RAPGEFL1 | Rap guanine nucleotide exchange factor (GEF)-like 1 |
| 3 | 31898109 | 31900366 | ENSCHAG00000015924 | SYT8 | synaptotagmin 8 |
| 3 | 31907006 | 31908504 | ENSCHAG00000015954 | novel | |
| 3 | 31913780 | 31915451 | ENSCHAG00000015975 | TNNI2A.1 | troponin I type 2a (skeletal, fast), tandem duplicate 1 |
| 3 | 31918862 | 31921460 | ENSCHAG00000016153 | TNNI2A.2 | troponin I type 2a (skeletal, fast), tandem duplicate 2 |
| 3 | 31928402 | 31935647 | ENSCHAG00000016160 | TNNI2 | troponin I2, fast skeletal type |
| 3 | 31942055 | 31943323 | ENSCHAG00000016191 | lncRNA | |
| 3 | 31958800 | 31981904 | ENSCHAG00000016193 | TNNI2 | troponin I2, fast skeletal type |
| 3 | 31963188 | 31964450 | ENSCHAG00000016209 | TNNI2 | troponin I2, fast skeletal type |
| 3 | 31968959 | 31974405 | ENSCHAG00000016273 | TNNI2 | troponin I, fast skeletal muscle-like |
| 3 | 31987347 | 31989147 | ENSCHAG00000016279 | TNNI2 | troponin I2, fast skeletal type |
| 3 | 31994267 | 31997377 | ENSCHAG00000016334 | TNNI2 | troponin I2, fast skeletal type |

| | | | | | |
|---|---|---|---|---|---|
| 3 | 32001722 | 32003702 | ENSCHAG00000016374 | TNNI2 | troponin I2, fast skeletal type |
| 3 | 32007386 | 32009098 | ENSCHAG00000016464 | TNNI2 | Clupea harengus troponin I, fast skeletal muscle-like (LOC105904758), mRNA. |
| 3 | 32012744 | 32013957 | ENSCHAG00000016508 | TNNI2 | troponin I2, fast skeletal type |
| 3 | 32021076 | 32028066 | ENSCHAG00000016547 | TNNI2A.4 | troponin I2, fast skeletal type |
| 3 | 32023986 | 32024535 | ENSCHAG00000016750 | TNNI2 | troponin I2, fast skeletal type |
| 3 | 32042563 | 32071473 | ENSCHAG00000016765 | CALDESMON | non-muscle caldesmon-like |
| 10 | 21564942 | 21565063 | ENSCHAG00000000386 | snoRNA | |
| 10 | 26242164 | 26247470 | ENSCHAG00000000387 | CCDC181 | coiled-coil domain containing 181 |
| 10 | 21565309 | 21565430 | ENSCHAG00000000401 | snoRNA | |
| 10 | 21566037 | 21566157 | ENSCHAG00000000418 | snoRNA | |
| 10 | 26247872 | 26251393 | ENSCHAG00000000436 | CPOX | coproporphyrinogen oxidase |
| 10 | 21566425 | 21566549 | ENSCHAG00000000440 | snoRNA | |
| 10 | 26255142 | 26258099 | ENSCHAG00000000469 | PRRG1 | proline rich and Gla domain 1 |
| 10 | 21566946 | 21567070 | ENSCHAG00000000514 | snoRNA | |
| 10 | 21592358 | 21595603 | ENSCHAG00000000664 | RASL11B | RAS-like, family 11, member B |
| 10 | 21598470 | 21745074 | ENSCHAG00000000822 | SCFD2 | sec1 family domain containing 2 |
| 10 | 26279154 | 26281406 | ENSCHAG00000001003 | ZBTB11 | Denticle herring_query56 |
| 10 | 26281625 | 26282282 | ENSCHAG00000001046 | ZBTB11 | Coho_salmon_query79 |
| 10 | 26282764 | 26283144 | ENSCHAG00000001063 | novel | |
| 10 | 26284353 | 26284448 | ENSCHAG00000001073 | SNORD14 | Small nucleolar RNA SNORD14 |
| 10 | 26284632 | 26284727 | ENSCHAG00000001080 | SNORD14 | Small nucleolar RNA SNORD14 |
| 10 | 26284931 | 26285026 | ENSCHAG00000001084 | SNORD14 | Small nucleolar RNA SNORD14 |
| 10 | 26285283 | 26285378 | ENSCHAG00000001094 | SNORD14 | Small nucleolar RNA SNORD14 |
| 10 | 26285574 | 26285669 | ENSCHAG00000001109 | SNORD14 | Small nucleolar RNA SNORD14 |

| | | | | | |
|---|---|---|---|---|---|
| 10 | 26285866 | 26285960 | ENSCHAG00000001120 | SNORD14 | Small nucleolar RNA SNORD14 |
| 10 | 26286235 | 26286330 | ENSCHAG00000001134 | SNORD14 | Small nucleolar RNA SNORD14 |
| 10 | 26286572 | 26286666 | ENSCHAG00000001146 | SNORD14 | Small nucleolar RNA SNORD14 |
| 10 | 26286869 | 26286963 | ENSCHAG00000001163 | SNORD14 | Small nucleolar RNA SNORD14 |
| 10 | 26287157 | 26287252 | ENSCHAG00000001174 | SNORD14 | Small nucleolar RNA SNORD14 |
| 10 | 26287516 | 26287610 | ENSCHAG00000001179 | SNORD14 | Small nucleolar RNA SNORD14 |
| 10 | 26287804 | 26287899 | ENSCHAG00000001185 | SNORD14 | Small nucleolar RNA SNORD14 |
| 10 | 26288155 | 26288249 | ENSCHAG00000001192 | SNORD14 | Small nucleolar RNA SNORD14 |
| 10 | 26288452 | 26288547 | ENSCHAG00000001197 | SNORD14 | Small nucleolar RNA SNORD14 |
| 10 | 26288817 | 26288912 | ENSCHAG00000001202 | SNORD14 | Small nucleolar RNA SNORD14 |
| 10 | 26289138 | 26289232 | ENSCHAG00000001212 | SNORD14 | Small nucleolar RNA SNORD14 |
| 10 | 22020303 | 22021975 | ENSCHAG00000001217 | GSX2 | GS homeobox 2 |
| 10 | 26289431 | 26289526 | ENSCHAG00000001220 | SNORD14 | Small nucleolar RNA SNORD14 |
| 10 | 26289723 | 26289818 | ENSCHAG00000001228 | SNORD14 | Small nucleolar RNA SNORD14 |
| 10 | 26289925 | 26291313 | ENSCHAG00000001243 | lncRNA | |
| 10 | 22026037 | 22026870 | ENSCHAG00000001249 | PDX1L | pancreas/duodenum homeobox protein 1-like |
| 10 | 26290078 | 26290173 | ENSCHAG00000001260 | SNORD14 | Small nucleolar RNA SNORD14 |
| 10 | 26290408 | 26290502 | ENSCHAG00000001267 | SNORD14 | Small nucleolar RNA SNORD14 |
| 10 | 26302540 | 26311445 | ENSCHAG00000001272 | DPT | dermatopontin |
| 10 | 26329511 | 26338856 | ENSCHAG00000001285 | ATP1B1A | ATPase Na+/K+ transporting subunit beta 1a |
| 10 | 22034602 | 22059625 | ENSCHAG00000001576 | PDGFRA | platelet-derived growth factor receptor, alpha polypeptide |
| 10 | 27476408 | 27523219 | ENSCHAG00000010108 | FNBP1L | formin binding protein 1 like |

| | | | | | |
|---|---|---|---|---|---|
| 10 | 27523130 | 27530702 | ENSCHAG00000010738 | SI:CH211-198N 5.11 | methylcrotonoyl-CoA carboxylase beta chain, mitochondrial-like |
| 10 | 27530650 | 27536911 | ENSCHAG00000010933 | RPE65B | retinoid isomerohydrolase RPE65 |
| 10 | 27537043 | 27545685 | ENSCHAG00000011503 | DEPDC1A | DEP domain containing 1 |
| 10 | 27553553 | 27685709 | ENSCHAG00000011575 | LRRC7 | leucine rich repeat containing 7 |
| 10 | 20494208 | 20548284 | ENSCHAG00000019366 | EPHB3 | EPH receptor B3 |
| 10 | 20831204 | 20852583 | ENSCHAG00000021082 | ALDH9A1A.1 | aldehyde dehydrogenase 9 family, member A1a, tandem duplicate 1 |
| 10 | 20852841 | 20857101 | ENSCHAG00000021442 | TMCO1 | transmembrane and coiled-coil domains 1 |
| 10 | 20857998 | 20868712 | ENSCHAG00000021499 | UCK2A | uridine-cytidine kinase |
| 10 | 20870941 | 20873588 | ENSCHAG00000021529 | MAGOH | protein mago nashi homolog |
| 10 | 20874429 | 20877972 | ENSCHAG00000021555 | CPT2 | carnitine palmitoyltransferase 2 |
| 10 | 20879109 | 20893305 | ENSCHAG00000021596 | PFAS | phosphoribosylformylglycinamidine synthase |
| 10 | 24999751 | 25030480 | ENSCHAG00000021680 | DIPK1A | divergent protein kinase domain 1A |
| 10 | 25035586 | 25039343 | ENSCHAG00000021694 | lncRNA | |
| 10 | 25039439 | 25045211 | ENSCHAG00000021695 | lncRNA | |
| 10 | 25053029 | 25066536 | ENSCHAG00000021706 | SCINLA | scinderin like a |
| 10 | 25066330 | 25070086 | ENSCHAG00000022110 | ZGC:109982 | zgc:109982 |
| 10 | 25072244 | 25074010 | ENSCHAG00000022113 | DR1 | down-regulator of transcription 1 |
| 10 | 25109758 | 25110771 | ENSCHAG00000022117 | lncRNA | |
| 10 | 25115655 | 25117298 | ENSCHAG00000022120 | GNG12 | si:dkey-44g17.6 |
| 10 | 25123881 | 25126178 | ENSCHAG00000022129 | GADD45AB | growth arrest and DNA damage inducible alpha |
| 10 | 25128212 | 25144250 | ENSCHAG00000022133 | SEC16B | SEC16 homolog B, endoplasmic reticulum export factor |
| 10 | 25156922 | 25158661 | ENSCHAG00000022182 | ZNF648 | zinc finger protein 648 |
| 10 | 25190558 | 25194017 | ENSCHAG00000022196 | GLULA | glutamate-ammonia ligase (glutamine synthase) a |
| 10 | 25199375 | 25211724 | ENSCHAG00000022250 | novel | |

| | | | | | |
|---|---|---|---|---|---|
| 10 | 25214741 | 25215979 | ENSCHAG00000022272 | lncRNA | |
| 10 | 25360248 | 25453127 | ENSCHAG00000022546 | SZT2 | SZT2 subunit of KICSTOR complex |
| 10 | 25667104 | 25840741 | ENSCHAG00000023174 | PTPRFA | protein tyrosine phosphatase receptor type Fa |
| 10 | 21073315 | 21122475 | ENSCHAG00000023224 | lncRNA | |
| 10 | 21074974 | 21123157 | ENSCHAG00000023263 | PSEUDOGENE | |
| 10 | 21105650 | 21109120 | ENSCHAG00000023281 | lncRNA | |
| 10 | 21129511 | 21137748 | ENSCHAG00000023283 | CBR4 | carbonyl reductase 4 |
| 10 | 21138281 | 21183110 | ENSCHAG00000023299 | PALLD | palladin, cytoskeletal associated protein |
| 10 | 21350609 | 21458637 | ENSCHAG00000023473 | FRYL | furry homolog, like |
| 10 | 21459206 | 21462560 | ENSCHAG00000024930 | OCIAD1 | OCIA domain containing 1 |
| 10 | 21463012 | 21468188 | ENSCHAG00000024945 | OCIAD2 | OCIA domain containing 2 |
| 10 | 21471199 | 21479834 | ENSCHAG00000024952 | DCUN1D4 | DCN1, defective in cullin neddylation 1, domain containing 4 (S. cerevisiae) |
| 10 | 21483386 | 21496359 | ENSCHAG00000024991 | novel | |
| 10 | 21497600 | 21501206 | ENSCHAG00000024997 | SGCB | sarcoglycan, beta (dystrophin-associated glycoprotein) |
| 10 | 21501708 | 21514660 | ENSCHAG00000025006 | SPATA18 | spermatogenesis associated 18 |
| 10 | 21548030 | 21563169 | ENSCHAG00000025013 | USP46 | ubiquitin specific peptidase 46 |
| 10 | 21564107 | 21567256 | ENSCHAG00000025041 | lncRNA | |
| 10 | 26172089 | 26181821 | ENSCHAG00000027469 | RORCA | nuclear receptor ROR-beta-like |
| 10 | 26188424 | 26207794 | ENSCHAG00000027470 | SH3GL1B | SH3 domain containing GRB2 like 1, endophilin A2 |
| 10 | 26211666 | 26216771 | ENSCHAG00000027478 | MPND | MPN domain containing |
| 10 | 26220359 | 26228506 | ENSCHAG00000027492 | STAP2B | signal transducing adaptor family member 2b |
| 10 | 26231207 | 26241775 | ENSCHAG00000027494 | FSD1 | fibronectin type III and SPRY domain containing 1 |
| 12 | 16045752 | 16047174 | ENSCHAG00000008006 | novel | |

| | | | | | |
|---|---|---|---|---|---|
| 12 | 16049213 | 16055058 | ENSCHAG00000008040 | lncRNA | |
| 12 | 16079212 | 16091737 | ENSCHAG00000008143 | MYHC | myosin heavy chain, fast skeletal muscle-like |
| 12 | 16124211 | 16125373 | ENSCHAG00000011128 | lncRNA | |
| 12 | 16138689 | 16150374 | ENSCHAG00000011333 | MYHC4 | cod_query41.83 |
| 12 | 16186421 | 16187545 | ENSCHAG00000019269 | novel | |
| 12 | 16223364 | 16235223 | ENSCHAG00000019283 | MYHZ2 | zebrafish_query90.68 |
| 12 | 16250204 | 16251937 | ENSCHAG00000019294 | HSPB15 | heat shock protein, alpha-crystallin-related, b15 |
| 12 | 16252644 | 16261948 | ENSCHAG00000019298 | SUSD2 | sushi domain containing 2 |
| 16 | 13955862 | 13983460 | ENSCHAG00000000936 | TULP3 | zmp:0000000711 |
| 16 | 13963775 | 13965116 | ENSCHAG00000001045 | novel | |
| 16 | 13983664 | 13988237 | ENSCHAG00000001076 | FOXM1 | forkhead box M1 |
| 16 | 13989283 | 13994724 | ENSCHAG00000001310 | SI:CH73-352P4.8 | si:ch73-352p4.8 |
| 16 | 13995965 | 13998293 | ENSCHAG00000003243 | PEX26 | peroxisomal biogenesis factor 26 |
| 16 | 14000020 | 14003844 | ENSCHAG00000003267 | USP18 | ubiquitin specific peptidase 18 |
| 16 | 14058043 | 14061495 | ENSCHAG00000003394 | FGF6B | fibroblast growth factor 6 |
| 16 | 14074810 | 14077952 | ENSCHAG00000003415 | RAD51AP1 | RAD51 associated protein 1 |
| 16 | 14081599 | 14088256 | ENSCHAG00000003478 | DYRK4 | dual specificity tyrosine phosphorylation regulated kinase 4 |
| 16 | 14092877 | 14100207 | ENSCHAG00000003806 | GALNT8 | polypeptide N-acetylgalactosaminyltransferase 8b, tandem duplicate 1 |
| 16 | 14122023 | 14200394 | ENSCHAG00000003909 | COG5 | component of oligomeric golgi complex 5 |
| 16 | 14185646 | 14187866 | ENSCHAG00000004711 | GPR22A | G protein-coupled receptor 22a |
| 16 | 14200176 | 14203385 | ENSCHAG00000004743 | DUS4L | dihydrouridine synthase 4 like |
| 16 | 14204120 | 14208631 | ENSCHAG00000004827 | BCAP29 | B cell receptor associated protein 29 |
| 16 | 14210452 | 14227276 | ENSCHAG00000004880 | SLCO1C1 | solute carrier organic anion transporter family, member 1C1 |

| | | | | | |
|---|---|---|---|---|---|
| 16 | 14244422 | 14308432 | ENSCHAG00000005408 | PDE3A | phosphodiesterase 3A, cGMP-inhibited |
| 16 | 14359704 | 14375859 | ENSCHAG00000005528 | AEBP2 | AE binding protein 2 |
| 16 | 14378135 | 14494204 | ENSCHAG00000005592 | PLEKHA5 | pleckstrin homology domain containing, family A member 5 |
| 16 | 14581273 | 14584783 | ENSCHAG00000006655 | novel | |
| 16 | 14587644 | 14614220 | ENSCHAG00000006661 | BICD1L | protein bicaudal D homolog 1-like |
| 16 | 14619789 | 14632823 | ENSCHAG00000006688 | FGD4B | FYVE, RhoGEF and PH domain containing 4b |
| 16 | 14634196 | 14638832 | ENSCHAG00000007917 | PRKAR2B | protein kinase cAMP-dependent type II regulatory subunit beta |
| 16 | 14642189 | 14649488 | ENSCHAG00000008000 | PIK3CG | phosphatidylinositol-4,5-bisphosphate 3-kinase catalytic subunit gamma |
| 16 | 14650825 | 14666577 | ENSCHAG00000008180 | NAMPT | si:dkey-145c18.3 |
| 16 | 14675217 | 14680960 | ENSCHAG00000009735 | ZNF800A | zinc finger protein 800a |
| 16 | 14684006 | 14827571 | ENSCHAG00000009841 | GRM8 | glutamate receptor, metabotropic 8a |
| 16 | 15115220 | 15154158 | ENSCHAG00000012375 | SEMA3C | sema domain, immunoglobulin domain (Ig), short basic domain, secreted, (semaphorin) 3C |
| 16 | 15157411 | 15165343 | ENSCHAG00000013260 | CD36 | CD36 molecule (thrombospondin receptor) |
| 16 | 15167547 | 15182850 | ENSCHAG00000013538 | GNAI1 | G protein subunit alpha i1 |
| 16 | 15220915 | 15433568 | ENSCHAG00000013742 | MAGI2A | membrane associated guanylate kinase, WW and PDZ domain containing 2a |
| 17 | 25255534 | 25272705 | ENSCHAG00000003278 | RREB1 | ras responsive element binding protein 1a |
| 17 | 25319456 | 25328701 | ENSCHAG00000003470 | LY86 | lymphocyte antigen 86 |
| 17 | 25337356 | 25346141 | ENSCHAG00000003517 | F13A1 | coagulation factor XIII A chain |
| 17 | 25359224 | 25369735 | ENSCHAG00000003862 | F13A1A.1 | goldfish_query45.03 |
| 17 | 25364646 | 25364759 | ENSCHAG00000004421 | U5 | U5 spliceosomal RNA |
| 17 | 18588033 | 18589772 | ENSCHAG00000009573 | FZD8A | frizzled-8-like |
| 17 | 18599026 | 18601014 | ENSCHAG00000009580 | | fucolectin-like |

| | | | | | |
|---|---|---|---|---|---|
| 17 | 18624901 | 18626636 | ENSCHAG00000009593 | novel | |
| 17 | 18659318 | 18662162 | ENSCHAG00000009606 | novel | |
| 17 | 18703198 | 18704757 | ENSCHAG00000009619 | CDH20 | Denticle herring_query68 |
| 17 | 18705596 | 18709014 | ENSCHAG00000009625 | novel | |
| 17 | 18714022 | 18733956 | ENSCHAG00000009643 | CDH20 | cadherin 20 |
| 17 | 18775519 | 18779502 | ENSCHAG00000010866 | lncRNA | |
| 19 | 6319922 | 6341388 | ENSCHAG00000002578 | GMPR | guanosine monophosphate reductase |
| 19 | 6344165 | 6477279 | ENSCHAG00000003500 | THRB | thyroid hormone receptor beta |

**Supplementary Table 5 -** Gene Ontology Enrichment analysis of genes in introgression regions (Table 1).

**Supplementary Table 6 -** Functions of single nucleotide polymorphisms within introgressed regions (Table1).

**Supplementary Table 7 -** Introgressed regions classified according to divergence and selection coefficient. Confident regions are ones where $d_{xy}$ between Baltic spring and White Sea (chosen as representative of Arctic Pacific herring) is below the 5% threshold and $d_{xy}$ between Baltic autumn and White Sea is above the 5% threshold. Very confident regions are ones where standard xpEHH is, in addition, below -2 and thus are under positive selection in Baltic spring-spawners. Other introgressed regions do not meet these criteria. Divergence is calculated using Baltic spring individuals who are homozygotes for Pacific herring alleles. As homozygosity varies by introgressed region, we calculated a $d_{xy}$ distribution per region. If there were no homozygous individuals, no $d_{xy}$ was calculated, as is the case of some regions in chromosome 16**.**

| Chromosome | Start | End | Divergence ($d_{xy}$) | | | | Selection | Confidence |
|---|---|---|---|---|---|---|---|---|
| | | | Baltic spring vs White sea | 5% $d_{xy}$ threshold | Baltic autumn vs White sea | 5% $d_{xy}$ threshold | Std xpEHH | |
| Chr 1 | 28920001 | 28960000 | 0.002 | 0.002 | 0.008 | 0.002 | 0.573 | |
| Chr 1 | 28960001 | 28980000 | 0.001 | 0.002 | 0.004 | 0.002 | -0.794 | Confident |
| Chr 1 | 28980001 | 29000000 | 0.001 | 0.002 | 0.004 | 0.002 | 0.291 | Confident |
| Chr 1 | 29000001 | 29020000 | 0.001 | 0.002 | 0.005 | 0.002 | 0.867 | Confident |
| Chr 1 | 29020001 | 29080000 | 0.003 | 0.002 | 0.005 | 0.002 | 0.886 | |
| Chr 3 | 31920001 | 31940000 | 0.002 | 0.001 | 0.006 | 0.002 | -1.894 | |
| Chr 3 | 31940001 | 31980000 | 0.002 | 0.001 | 0.007 | 0.002 | 1.488 | |
| Chr 3 | 31980001 | 32000000 | 0.003 | 0.001 | 0.008 | 0.002 | 2.294 | |
| Chr 3 | 32000001 | 32020000 | 0.002 | 0.001 | 0.006 | 0.002 | -0.835 | |
| Chr 3 | 32020001 | 32040000 | 0.002 | 0.001 | 0.007 | 0.002 | -1.678 | |
| Chr 10 | 20540001 | 20600000 | 0.003 | 0.001 | 0.006 | 0.002 | 0.667 | |
| Chr 10 | 20820001 | 20860000 | 0.003 | 0.001 | 0.012 | 0.002 | 0.960 | |
| Chr 10 | 21120001 | 21140000 | 0.000 | 0.001 | 0.003 | 0.002 | -1.235 | Confident |
| Chr 10 | 21380001 | 21400000 | 0.001 | 0.001 | 0.007 | 0.002 | -2.274 | Very Confident |
| Chr 10 | 21480001 | 21500000 | 0.001 | 0.001 | 0.004 | 0.002 | -0.339 | Confident |
| Chr 10 | 21500001 | 21580000 | 0.001 | 0.001 | 0.006 | 0.002 | -1.125 | Confident |
| Chr 10 | 21580001 | 21600000 | 0.002 | 0.001 | 0.006 | 0.002 | -2.322 | |
| Chr 10 | 21940001 | 21960000 | 0.002 | 0.001 | 0.004 | 0.002 | -0.478 | |
| Chr 10 | 22040001 | 22060000 | 0.001 | 0.001 | 0.005 | 0.002 | -0.671 | Confident |
| Chr 10 | 25040001 | 25060000 | 0.004 | 0.001 | 0.005 | 0.002 | -1.533 | |
| Chr 10 | 25060001 | 25080000 | 0.002 | 0.001 | 0.003 | 0.002 | -2.096 | |
| Chr 10 | 25080001 | 25100000 | 0.002 | 0.001 | 0.004 | 0.002 | -2.088 | |
| Chr 10 | 25100001 | 25120000 | 0.000 | 0.001 | 0.003 | 0.002 | -3.154 | Very Confident |

| | | | | | | | | |
|---|---|---|---|---|---|---|---|---|
| Chr 10 | 25120001 | 25140000 | 0.001 | 0.001 | 0.003 | 0.002 | -3.094 | Very Confident |
| Chr 10 | 25140001 | 25160000 | 0.001 | 0.001 | 0.006 | 0.002 | -3.943 | Very Confident |
| Chr 10 | 25160001 | 25180000 | 0.001 | 0.001 | 0.004 | 0.002 | -4.061 | Very Confident |
| Chr 10 | 25180001 | 25200000 | 0.001 | 0.001 | 0.005 | 0.002 | -3.525 | Very Confident |
| Chr 10 | 25200001 | 25220000 | 0.001 | 0.001 | 0.004 | 0.002 | -2.812 | Very Confident |
| Chr 10 | 25380001 | 25400000 | 0.003 | 0.001 | 0.005 | 0.002 | 1.032 | |
| Chr 10 | 25680001 | 25700000 | 0.004 | 0.001 | 0.004 | 0.002 | -0.112 | |
| Chr 10 | 26200001 | 26240000 | 0.003 | 0.001 | 0.005 | 0.002 | -1.787 | |
| Chr 10 | 26300001 | 26320000 | 0.003 | 0.001 | 0.005 | 0.002 | -2.419 | |
| Chr 10 | 26320001 | 26340000 | 0.001 | 0.001 | 0.002 | 0.002 | -2.859 | Very Confident |
| Chr 10 | 26340001 | 26360000 | 0.001 | 0.001 | 0.003 | 0.002 | -3.723 | |
| Chr 10 | 27520001 | 27560000 | 0.002 | 0.001 | 0.007 | 0.002 | 0.500 | |
| Chr 12 | 16060001 | 16080000 | 0.002 | 0.002 | 0.005 | 0.002 | 1.092 | |
| Chr 12 | 16140001 | 16160000 | 0.001 | 0.002 | 0.004 | 0.002 | 1.650 | |
| Chr 12 | 16180001 | 16200000 | 0.000 | 0.002 | 0.003 | 0.002 | 1.137 | |
| Chr 12 | 16200001 | 16220000 | 0.001 | 0.002 | 0.004 | 0.002 | 1.003 | |
| Chr 12 | 16220001 | 16240000 | 0.001 | 0.002 | 0.003 | 0.002 | 0.739 | Confident |
| Chr 16 | 13980001 | 14000000 | 0.000 | 0.002 | 0.003 | 0.002 | -1.279 | Confident |
| Chr 16 | 14080001 | 14100000 | 0.001 | 0.001 | 0.002 | 0.002 | -1.139 | Confident |
| Chr 16 | 14100001 | 14120000 | NA | 0.001 | NA | 0.002 | 0.270 | |
| Chr 16 | 14120001 | 14240000 | 0.002 | 0.002 | 0.004 | 0.002 | -1.442 | |
| Chr 16 | 14380001 | 14420000 | 0.001 | 0.002 | 0.003 | 0.002 | -0.757 | Confident |
| Chr 16 | 14420001 | 14440000 | 0.001 | 0.002 | 0.004 | 0.002 | -1.612 | Confident |
| Chr 16 | 14440001 | 14460000 | 0.001 | 0.002 | 0.003 | 0.002 | -1.303 | Confident |
| Chr 16 | 14460001 | 14500000 | 0.001 | 0.002 | 0.003 | 0.002 | -1.603 | Confident |
| Chr 16 | 14500001 | 14520000 | 0.002 | 0.002 | 0.009 | 0.002 | -1.452 | |
| Chr 16 | 14520001 | 14580000 | 0.002 | 0.002 | 0.005 | 0.002 | -1.727 | Confident |
| Chr 16 | 14580001 | 14600000 | 0.001 | 0.002 | 0.004 | 0.002 | -2.192 | Very Confident |
| Chr 16 | 14600001 | 14640000 | 0.001 | 0.002 | 0.004 | 0.002 | -2.639 | Very Confident |

| | | | | | | | | |
|---|---|---|---|---|---|---|---|---|
| Chr 16 | 14640001 | 14700000 | 0.001 | 0.002 | 0.003 | 0.002 | -2.508 | Very Confident |
| Chr 16 | 14700001 | 14720000 | 0.001 | 0.002 | 0.004 | 0.002 | -1.382 | Confident |
| Chr 16 | 14720001 | 14740000 | NA | NA | NA | NA | NA | |
| Chr 16 | 14740001 | 14760000 | NA | NA | NA | NA | NA | |
| Chr 16 | 15140001 | 15180000 | NA | NA | NA | NA | NA | |
| Chr 16 | 15180001 | 15220000 | NA | NA | NA | NA | NA | |
| Chr 16 | 15220001 | 15240000 | NA | NA | NA | NA | NA | |
| Chr 16 | 15240001 | 15260000 | 0.002 | 0.002 | 0.006 | 0.002 | -0.917 | |
| Chr 16 | 15320001 | 15340000 | 0.001 | 0.002 | 0.007 | 0.002 | -2.209 | Very Confident |
| Chr 16 | 15340001 | 15360000 | 0.001 | 0.002 | 0.004 | 0.002 | -2.351 | Very Confident |
| Chr 17 | 18540001 | 18600000 | 0.001 | 0.001 | 0.003 | 0.002 | -4.369 | |
| Chr 17 | 18600001 | 18700000 | 0.002 | 0.001 | 0.008 | 0.002 | -4.067 | |
| Chr 17 | 18700001 | 18720000 | 0.006 | 0.001 | 0.009 | 0.002 | -3.264 | |
| Chr 17 | 18720001 | 18740000 | 0.002 | 0.001 | 0.003 | 0.002 | -4.078 | |
| Chr 17 | 18740001 | 18760000 | 0.001 | 0.001 | 0.004 | 0.002 | -2.870 | |
| Chr 17 | 18760001 | 18780000 | 0.002 | 0.001 | 0.004 | 0.002 | -1.957 | |
| Chr 17 | 25280001 | 25300000 | 0.002 | 0.001 | 0.005 | 0.002 | 0.516 | |
| Chr 17 | 25300001 | 25320000 | 0.004 | 0.001 | 0.004 | 0.002 | 1.769 | |
| Chr 17 | 25340001 | 25360000 | 0.003 | 0.001 | 0.003 | 0.002 | 1.803 | |
| Chr 19 | 6360001 | 6380000 | 0.001 | 0.002 | 0.006 | 0.002 | -2.987 | Very Confident |

**Supplementary Table 8 -** Blastn alignment results between the 1st exon of the retina specific THRB isoform of zebrafish and the ENSEMBL 114 Atlantic herring annotation.

| Genomic Location | Overlapping Gene(s) | Orientation | Query start | Query end | Query ori | Length | Score | E-val | %ID |
|---|---|---|---|---|---|---|---|---|---|
| 19:6359064-6359164 | THRB | Reverse | 157 | 257 | Forward | 101 | 65.7 | 3.28E-09 | 83.2 |
| 14:5541186-5541205 | ENSCHAG00020016078 | Forward | 43 | 62 | Forward | 20 | 40 | 0.18 | 100 |
| 3:19906204-19906223 | HSF4 | Forward | 109 | 128 | Forward | 20 | 40 | 0.18 | 100 |
| 22:546373-546391 | | Reverse | 239 | 257 | Forward | 19 | 38.1 | 0.7 | 100 |
| 1:9087062-9087080 | | Forward | 246 | 264 | Forward | 19 | 38.1 | 0.7 | 100 |
| 3:21561243-21561260 | KCNQ1.2 | Reverse | 137 | 154 | Forward | 18 | 36.1 | 2.8 | 100 |
| 23:10770322-10770339 | FAM20CB | Forward | 37 | 54 | Forward | 18 | 36.1 | 2.8 | 100 |
| 17:1154660-1154681 | LAMA1 | Reverse | 97 | 118 | Forward | 22 | 36.1 | 2.8 | 95.5 |
| 11:15465263-15465280 | | Reverse | 98 | 115 | Forward | 18 | 36.1 | 2.8 | 100 |
| 10:5836176-5836193 | | Forward | 30 | 47 | Forward | 18 | 36.1 | 2.8 | 100 |
| 10:26100868-26100885 | SEMA6BB | Forward | 226 | 243 | Forward | 18 | 36.1 | 2.8 | 100 |
| 5:1742047-1742064 | | Reverse | 98 | 115 | Forward | 18 | 36.1 | 2.8 | 100 |

**Supplementary Table S9** - List of primers used for genotyping and gene expression analyses

| Gene | Primer | Sequence (5′–3′) | Product size (bp) |
|---|---|---|---|
| THRb | Forward | TGCCCGACATGTATGAGATGC | 402 |
| THRb | Reverse | CGCTCATACCAGAGAAAGGTCC | |
| Actin | Forward | CACCATTGGAAACGAGAGGT | 147 |
| Actin | Reverse | GTGTTGGCGTACAGGTCCTT | |
| EF1A | Forward | TTATCGGCCACGTCGACTC | 219 |
| EF1A | Reverse | ACTTGCCGGTCTCAAACTT | |
| LSW1 | Forward | AAATTCTGTTTACGCTGCCAGAA | 158 |
| LSW1 | Reverse | TAAACATCCACACTGTGGCAAGG | |
| LSW2 | Forward | CGAAGCGGCTTTTGCTGC | 130 |
| LSW2 | Reverse | GATCCACCGTGGCGCAATGT | |

**Supplementary Table 10** - Differential expression of LSW1 and LSW2 in homozygous and heterozygous THRB genotypes based on qPCR normalization with ACTIN and EF1A.
